# Androgen deprivation upregulates SPINK1 expression and potentiates cellular plasticity in prostate cancer

**DOI:** 10.1101/562652

**Authors:** Ritika Tiwari, Nishat Manzar, Vipul Bhatia, Anjali Yadav, Shannon Carskadon, Nilesh Gupta, Amina Zoubeidi, Nallasivam Palanisamy, Bushra Ateeq

## Abstract

The Serine Peptidase Inhibitor, Kazal type 1 (SPINK1) overexpression represents ~10-25% of the prostate cancer (PCa) cases associated with shorter recurrence-free survival and poor prognosis. Nonetheless, androgen-deprivation therapy (ADT) remains the mainstay treatment for locally advanced and metastatic PCa patients. However, majority of these individuals eventually progress to castration-resistant stage, and a subset of these patients develop ADT-induced neuroendocrine PCa. Despite adverse effects of ADT, possible role of androgen signaling in SPINK1-mediated prostate oncogenesis remains unexplored. Here, we show that androgen receptor (AR) and its corepressor, the RE1-silencing transcription factor (REST), occupy *SPINK1* promoter and functions as a direct transcriptional repressor of *SPINK1*, thus blocking AR signaling via ADT relieves its repression, leading to SPINK1 upregulation. In agreement, an inverse association between SPINK1 levels and AR expression was observed across multiple PCa cohorts, and in neuroendocrine differentiated cells. While, lineage reprogramming factor SOX2 in turn binds to *SPINK1* promoter leading to its transactivation in androgen-deprived conditions with concomitant increase in neuroendocrine markers. Additionally, we also confirm the role of *SPINK1* in epithelial-mesenchymal transition, drug resistance, stemness and cellular plasticity. Moreover, we show that Casein Kinase 1 inhibitor stabilizes the REST levels, which in cooperation with AR, conjures transcriptional repression of *SPINK1* expression, and impedes SPINK1-mediated oncogenesis. Collectively, our findings provide a plausible explanation to the paradoxical clinical outcomes of ADT, possibly due to increased SPINK1 levels. This study highlights the need to take a well-informed decision prior to ADT and develop alternative therapeutic strategies for castrate-resistant PCa patients.

## Introduction

Genetic rearrangement involving androgen-driven promoter region of the serine protease gene, *TMPRSS2* and the coding region of *ERG*, a member of *ETS* (E26 transformation-specific) transcription factor family occurs in almost half of the prostate cancer (PCa) cases (Tomlins et al, 2005). Subsequently, with the technological advances in genomics, numerous other molecular subtypes such as, fusion involving other members of *ETS* family, (*ETV1*, *ETV4*, *FLI1* and *NDRG1*); *RAF* kinase rearrangements; *SPOP/CHD1* alterations; mutations in *FOXA1* and *IDH1* have also been discovered (Abeshouse et al, 2015; Tomlins et al, 2007; Tomlins et al, 2006). While *TMPRSS2-ERG* fusion is the most common subtype, overexpression of *SPINK1* constitutes a substantial ~10-25% of the total PCa cases exclusively in *ETS*-fusion negative subtype (Huang et al, 2016; Tomlins et al, 2008). However, several independent studies show expression of SPINK1 and ERG in two distinct foci within a prostate gland, indicating that these two events are either independent or SPINK1 overexpression to be a sub-clonal event after *TMPRSS2-ERG* genetic rearrangement (Brooks et al, 2015; Flavin et al, 2014; Huang et al, 2016). Notably, *SPINK1*-positive patients show rapid progression to biochemical recurrence as compared to *ETS*-fusion positive (Leinonen et al, 2010; Tomlins et al, 2008). In a Finnish PCa cohort, SPINK1-positive patients exhibit an association with an early progression to castration resistance (Leinonen et al, 2010). Further, intermediate or high risk localized PCa patients who endured radical prostatectomy show positive correlation between SPINK1 and biochemical recurrence subsequently leading to disease-specific mortality (Johnson et al, 2016).

Androgen receptor (AR) signaling axis plays critical role in the PCa pathogenesis and progression. Hence, androgen deprivation therapy (ADT) remains the basis for PCa treatment, however the disease often relapses to an androgen-independent advanced stage known as castrate-resistant prostate cancer (CRPC), often associated with poor patient prognosis (Karantanos et al, 2013; Lonergan & Tindall, 2011; Sun et al, 2012b). Numerous mechanisms that restore androgen signaling in CRPC individuals have been proposed, such as mutations in the AR ligand binding domain (F877L and T878A), constitutively active variants of AR (AR-V7 and ARv567es), AR amplification, or steroid-inducible glucocorticoid receptor that activates AR target genes (Antonarakis et al, 2014; Arora et al, 2013; Chen et al, 2004; Taplin et al, 1995), suggesting that these alteration in *AR* or AR-signaling ensue as an adaptive response to ADT. Current treatment regimen for CRPC patients include FDA-approved second generation anti-androgens such as enzalutamide or MDV3100 (blocks the nuclear translocation of AR or binding to its genomic sites) and abiraterone acetate (irreversible steroidal CYP17A1 inhibitor, targets adrenal or intratumoral androgen biosynthesis) (Attard et al, 2005; de Bono et al, 2011; Scher et al, 2010; Tran et al, 2009). Although, these anti-androgens are known to prolong the overall survival of PCa patients, but the response is temporary, and the disease eventually progresses. A subset of CRPC patients (~20% of advanced drug-resistant cases) elude selective pressure of ADT by minimizing its dependency on the AR signaling are identified as neuroendocrine (NE) PCa, often associated with poor prognosis and patient outcome (Rickman et al, 2017). NEPC exhibits a distinct phenotype characterized by reduced or no expression of AR and AR-regulated genes, and increased expression of NEPC markers such as Synaptophysin (SYP), Chromogranin A (CHGA), and Enolase 2 (ENO2) (Beltran et al, 2014). Several molecular mechanisms have been ascribed to the transdifferentiation of CRPC to NEPC, for instance, frequent genetic alterations involving the *TP53* and the retinoblastoma-1-encoding gene *RB1* are associated with poorly differentiated NEPC tumors (Beltran et al, 2016; Tan et al, 2014). Moreover, N-Myc amplification, BRN2 upregulation, mitotic deregulation via Aurora kinase A (AURKA), alternative splicing by serine/arginine repetitive matrix4 (SRRM4), and loss of repressor element-1 (RE-1) silencing transcription factor (REST) expression are known to have a role in NE transdifferentiation (Beltran et al, 2011; Bishop et al, 2017; Dardenne et al, 2016; Li et al, 2017; Svensson et al, 2014). Mounting evidence suggests that REST expression is downregulated in relapsed PCa (Svensson et al, 2014), which has also been attributed to NE differentiation of PCa cells (Lin et al, 2016; Svensson et al, 2014; Zhu et al, 2014). Interestingly, REST expression is positively regulated by androgen signaling, and it serves as a transcriptional co-regulator of AR (Li et al, 2017; Svensson et al, 2014).

Although, SPINK1 overexpression has been associated with rapid progression to biochemical recurrence and aggressive stage of PCa, nonetheless the regulatory mechanism involved in SPINK1 upregulation and its functional significance in the advancement of disease is largely undefined. In this study we have explored the mechanism involved in SPINK1 overexpression and its functional implication in PCa tumorigenesis. Importantly, we show that AR antagonists unexpectedly have a positive effect on the transcriptional regulation of *SPINK1*, and consecutive increase in SPINK1 level, further positively associates with NE-phenotype. Our findings demonstrate that *SPINK1* is transcriptionally suppressed by AR and its co-repressor REST, a negative master regulator of neurogenesis, and suggests a possible role of SPINK1 in NEPC transdifferentiation. Collectively, our findings alert against the widespread application of potent AR antagonist used for PCa treatment, and highlight expected clinical complication associated with increased expression of SPINK1 as well as therapy-induced NEPC.

## Results

### Expression of SPINK1 and Androgen Receptor is inversely correlated in prostate cancer patients

Altered AR signaling and AR-binding profile has been extensively studied in localized PCa and CRPC (Sharma et al, 2013). It has been known that AR binds with other cofactors, such as GATA2, octamer transcription factor 1 (Oct1), Forkhead box A1 (FoxA1) and nuclear factor 1 (NF-1) to mediate cooperative transcriptional activity of the target genes (Wang et al, 2007). Thus, we sought to discover the possible link between *SPINK1* and *AR* expression in PCa patients, and stratified patients available at TCGA-PRAD (The Cancer Genome Atlas Prostate Adenocarcinoma) cohort based on high and low expression of *AR*. The patients with higher expression of *AR* showed a significantly lower expression of *SPINK1* and contrariwise (Fig 1A). To further confirm this association, we performed immunohistochemical (IHC) analysis for the expression of SPINK1 and AR on tissue microarrays (TMA) comprising of PCa patient specimens (n=237). Important to note that all of these cases underwent radical prostatectomy without any hormone or radiation therapy. In concordance with TCGA data analysis, our IHC findings reveal that SPINK1-positive patients exhibit low or negative staining for AR expression, while SPINK1-negative patients show higher or medium AR staining (Fig 1B and Supplementary Fig S1A). Importantly, about ~67% of the SPINK1+ patients (34 out of 51) demonstrate either low or negative staining for AR expression (Fisher’s exact test, *P*<0.0004) (Fig 1C and D). Based on our findings, we conjecture that *SPINK1* is one of the AR repressed genes, hence we next examined the expression of other members of AR repressor complex (*NCOR1*, *NCOR2* and *NRIP1*) including AR using TCGA-PRAD cohort, and the patients were sorted based on *SPINK1* high and low expression by employing quartile-based normalization (Dillies et al, 2013). Interestingly, we found that *SPINK1* expression is also negatively associated with other AR repressive complex members (Supplementary Fig S1B). Additionally, we investigated the correlation of *SPINK1* and AR signaling score using transcriptomic data from two independent PCa cohorts, Memorial Sloan Kettering Cancer Center (MSKCC) and TCGA-PRAD. As expected, lower AR signaling score in *SPINK1*-positive patients was recorded as compared to the *SPINK1*-negative patients (Supplementary Fig S1C). Taken together, our findings show an inverse association between *SPINK1* expression and AR signaling in PCa patients, indicating that overexpression of SPINK1 in *SPINK1*-postive subtype is owing to the loss of AR-mediated repression during prostate cancer progression.

**Figure 1.**
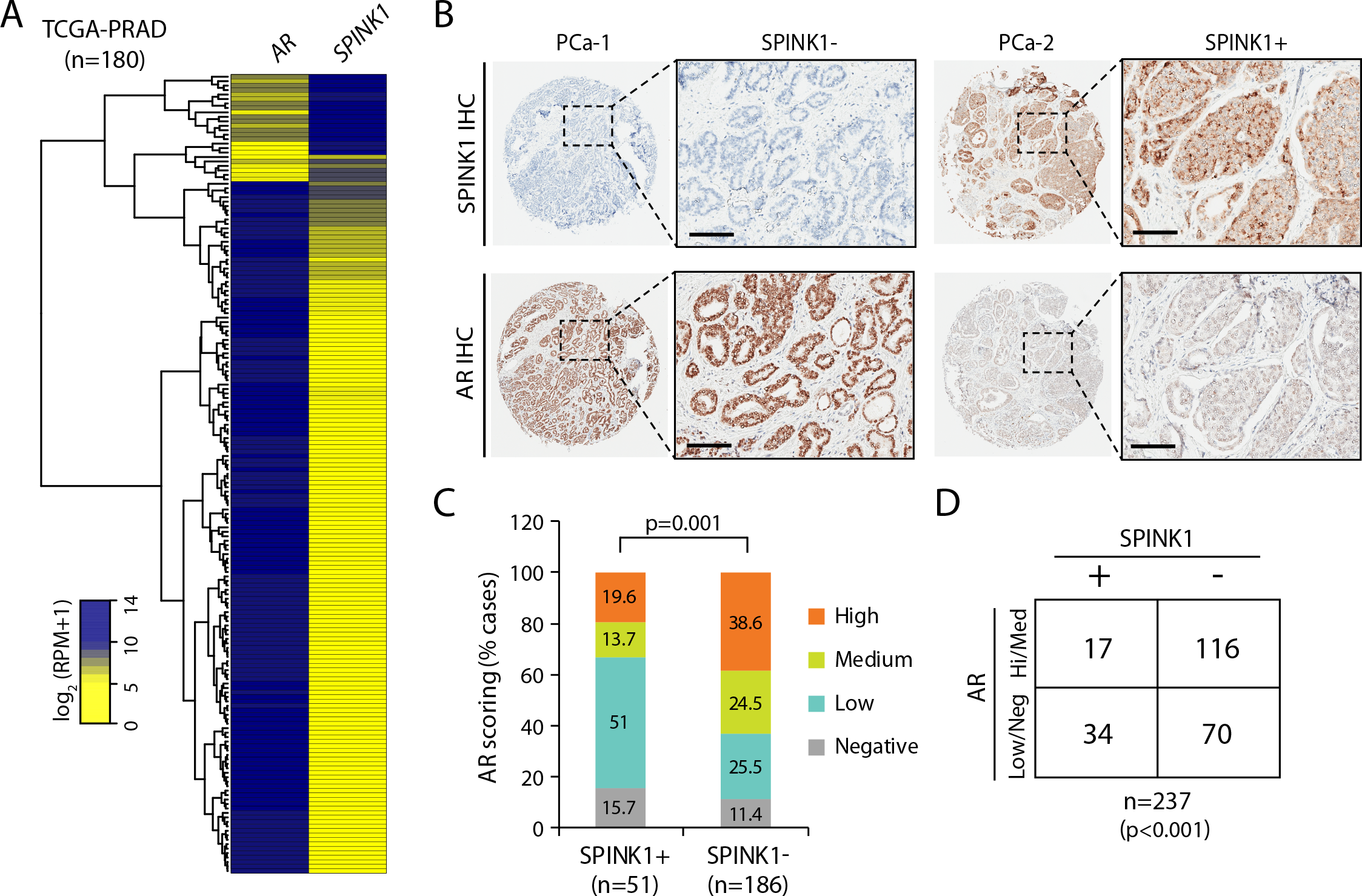
SPINK1 is negatively correlated with AR in PCa patient cohorts. (A) Heatmap depicting *AR* and *SPINK1* expression in TCGA-PRAD cohort (n=180). Shades of yellow and blue represents expression values in log_2_ (RPM+1). (B) Representative micrographs depicting PCa tissue microarray (TMA) cores (n=237), immunostained for SPINK1 and AR expression by immunohistochemistry (IHC). Panel on the top show representative IHC for SPINK1 in SPINK1-negative (SPINK1−) and SPINK1-positive (SPINK1+) patients. Bottom panel represents IHC for AR expression in same patients. Scale bar represents 500 μm and 100 μm for the entire core and the inset image respectively. (C) Bar plot showing percentage of IHC scoring for AR in the SPINK1− and SPINK1+ patients’ specimens. *P*-value for the Chi-Square test is indicated. (D) Contingency table for the AR and SPINK1 status. Patients showing high or medium expression of AR were grouped as AR-(Hi/Med), while patients with low or null AR expression were indicated as AR-(Low/Neg). *P*-value for Fisher’s exact test is indicated.

### AR antagonists trigger SPINK1 upregulation by relieving AR-mediated repression

Since, an inverse association between *SPINK1* expression and *AR* signaling was observed in three independent PCa cohorts (TCGA-PRAD, MSKCC) including ours (Fig 1), thus we examined the role of active AR signaling in the regulation of SPINK1 using PCa cell lines, 22RV1 (endogenously SPINK1-positive) and androgen responsive VCaP cells (*TMPRSS2-ERG* fusion positive) (Supplementary Fig S2A). Stimulating 22RV1 cells with synthetic androgen, R1881 (10nM) (Jin et al, 2007), results in a significant decrease in expression of *SPINK1* with a concomitant increase in the expression of AR target genes such as *KLK3*, *TMPRSS2* and *FKBP5* (Fig 2A-C and Supplementary Fig S2C). To further investigate whether similar effect on SPINK1 expression could be rendered by sub-physiological concentration of androgen, 22RV1 cells were stimulated with remarkably much lower concentrations of R1881 (0.01 and 0.1nM), interestingly both ~0.1nM and 1nM of R1881 were equally efficacious in repressing the expression of *SPINK1* transcript (Supplementary Fig S2B). Similarly, VCaP cells stimulated with R1881 (10nM) show a significant decline in the expression of *SPINK1* both at transcript and protein levels, while an increase in the expression of AR target genes (*KLK3*, *TMPRSS2* and *FKBP5*) was noticed (Fig 2D-F and Supplementary Fig S2C). Further, we also analyzed the publicly available datasets (GSE71797 and GSE51872), wherein 22RV1 and VCaP cells stimulated with R1881 and dihydrotestosterone (DHT) respectively, exhibits reduced expression of several previously known AR repressed genes, namely *DDC*, *OPRK1*, *NOV* and *SERPIN1* (Wu et al, 2014; Zhao et al, 2012) besides *SPINK1* (Fig 2G and H).

**Figure 2.**
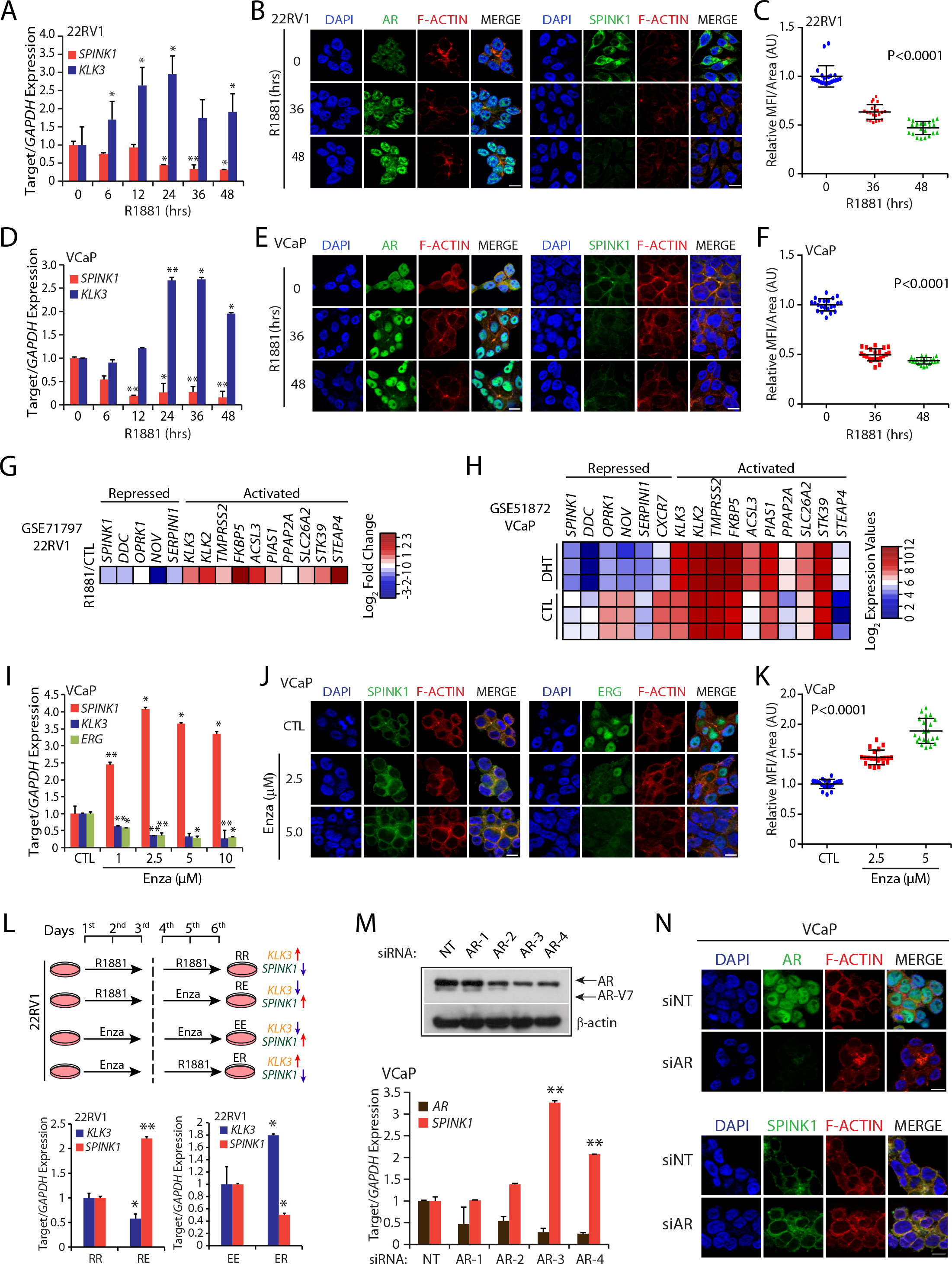
Androgen signaling negatively regulates *SPINK1* expression in prostate cancer. (A) Quantitative PCR data showing relative expression of *SPINK1* and *KLK3* in 22RV1 (*SPINK1+*) cells upon androgen (R1881) stimulation (10nM) at the indicated timepoints. (B) Immunostaining for SPINK1 and AR expression in 22RV1 (*SPINK1+*) cells upon androgen stimulation (10nM R1881) at different timepoints. F-actin and nucleus was stained by TRITC-phalloidin and DAPI respectively. Scale bar represents 10μm. (C) Same as (B), except quantification of immunofluorescence images. Dot plot represents relative mean fluorescence intensity per unit μm^2^ (in arbitrary units; A.U.). (D) Same as in (A), except VCaP cells (*TMPRSS2-ERG+*) were used. (E) Same as in (B), except VCaP cells were used. (F) Same as in (C), except quantification of VCaP cells. (G) Heatmap depicting relative expression of androgen regulated genes in 22RV1 cells upon androgen stimulation (GSE71797). Shades of red and blue represent log_2_ fold change from FPKM values of RNA-Seq data. (H) Same as in (G), except VCaP cells (GSE51872). Shades of red and blue represent log_2_ gene expression values. (I) Quantitative PCR data showing relative expression of *SPINK1*, *KLK3* and *ERG* in VCaP cells treated with enzalutamide at the indicated concentrations. (J) Immunostaining for SPINK1 and ERG using same cells as in (I). F-actin and nucleus was stained by TRITC-phalloidin and DAPI respectively. Scale bar represents 10μm. (K) Same as (J), except quantification of immunofluorescence images. Dot plot represents relative mean fluorescence intensity (MFI) per μm^2^ (in arbitrary units; A.U.) of enzalutamide treated VCaP cells. (L) Schema showing sequential treatment of 22RV1 cells with synthetic androgen R1881 (10nM) for 3 days followed by enzalutamide (10μM) for 3 days. The order of R1881 and enzalutamide was reversed in the second set of experiment keeping other conditions identical. Quantitative PCR data showing relative expression of *KLK3* and *SPINK1* in 22RV1 cells sequentially treated with R1881 or enzalutamide (bottom panel). (M) Quantitative PCR data showing relative expression of *AR* and *SPINK1* in siRNA mediated AR silenced VCaP cells with respect control. Immunoblot for AR levels in *AR* silenced VCaP cells. β-actin was used as a loading control. (N) Immunostaining for AR and SPINK1 using same cells as in (M). F-actin and nucleus was stained by TRITC-phalloidin and DAPI respectively. Scale bar represents 10μm. Statistical significance was calculated by one-way ANOVA with Tukey’s post hoc test for multiple comparisons for the panel (C) (F) and (K). In all panels except, (B), (E), (G), (H) (J) and (N), biologically independent samples were used (n=3); data represents mean ± SEM. \**P*≤ 0.05 and \*\**P*≤ ≤ 1.1 using two-tailed unpaired Student’s *t* test.

Non-steroidal pharmacological inhibitors for AR, namely bicalutamide (Bic) and enzalutamide (Enza) have been widely used for the treatment of locally advanced non-metastatic as well as metastatic prostate cancer (Chen et al, 2009; Scher et al, 2010; Tran et al, 2009), therefore we determined the effect of these anti-androgens on the expression of *SPINK1.* To antagonize AR signaling, VCaP cells were treated with Enza and SPINK1 expression was examined, notably a remarkable increase in the *SPINK1* both at transcript (~4-fold) and protein levels was observed, accompanied with reduced expression of androgen driven-genes such as *KLK3* and *ERG* (Fig 2I-K). To further corroborate these findings, we treated VCaP cells with Bic (25 and 50μM) and found a significant increase in the SPINK1 expression (Supplementary Fig S2D-F). Next, to evaluate any change in the oncogenic properties of the R1881-stimulated and/or Bic or Enza treated VCaP cells, Transwell migration assay was performed. A significant increase in the migratory properties of the androgen-stimulated VCaP cells treated with Bic or Enza was observed (Supplementary Fig S2G). Notably, 22RV1 cells are not much responsive to androgen as VCaP cells, therefore we used a strategy to modulate the AR signaling via priming 22RV1 cells either with R1881 or Enza for 3 days, followed by Enza treatment or R1881 stimulation for the next 3 consecutive days. As anticipated, blocking androgen signaling with Enza in the androgen-primed 22RV1 cells result in significant increase in *SPINK1* expression, while Enza-treated 22RV1 cells stimulated with R1881 show a significant repression of *SPINK1* transcript (Fig 2L). To examine the effect of long-term DHT treatment on *SPINK1* expression, 22RV1 cells were cultured in DHT (8nM) for 2 months, which resulted in more than ~80% reduction in *SPINK1* expression (Supplementary Fig S2H). Conversely, long-term blockade of androgen signaling in 22RV1 cells using Bic (5μM) led to significant increase (~1.5 folds) in the *SPINK1* expression (Supplementary Fig S2I). Similar results were obtained in androgen-regulated CWR22Pc cells, a derivative cell line of CWR22 xenograft subjected to long-term Bic treatment (Supplementary Fig S2J).

Alternative to pharmacological inhibition of AR signaling, we used small interfering RNA (siRNA) approach to abolish AR expression in 22RV1 and VCaP cells and examine any change in SPINK1 levels. Similar to the small molecule inhibition of AR signaling, siRNA-mediated *AR*-silenced 22RV1 cells exhibit moderate increase in the expression of *SPINK1* (Supplementary Fig S2L-N), while a robust increase (~3-fold) in the SPINK1 transcript and protein was observed in *AR*-silenced VCaP cells (Fig 2M and N and Supplementary Fig S2K). Taken together, our findings demonstrate that AR signaling negatively regulates SPINK1 expression and draws attention to AR antagonists mediated upregulation of SPINK1 in prostate cancer.

### AR directly binds to *SPINK1* promoter and regulates its expression

The role of AR has been extensively characterized both as a global transcriptional activator as well as repressor (Hu & Lazar, 2000; Zhao et al, 2012). To examine whether AR directly regulates *SPINK1* transcription we looked for putative AR binding sites in the *SPINK1* promoter region, and scanned the *SPINK1* promoter for the presence of androgen response elements (AREs) by employing publicly available transcription factor binding prediction software, JASPAR (Khan et al, 2018) and MatInspector (Cartharius et al, 2005). Several putative AREs within the ~5kb region upstream of transcription start site (TSS) of the *SPINK1* gene were identified (Fig 3A). Further, analysis of the publicly available Chromatin Immunoprecipitation-Sequencing (ChIP-Seq) dataset for AR binding in androgen stimulated VCaP cells (GSE8428) revealed another putative ARE on the *SPINK1* promoter (Fig 3B).

**Figure 3.**
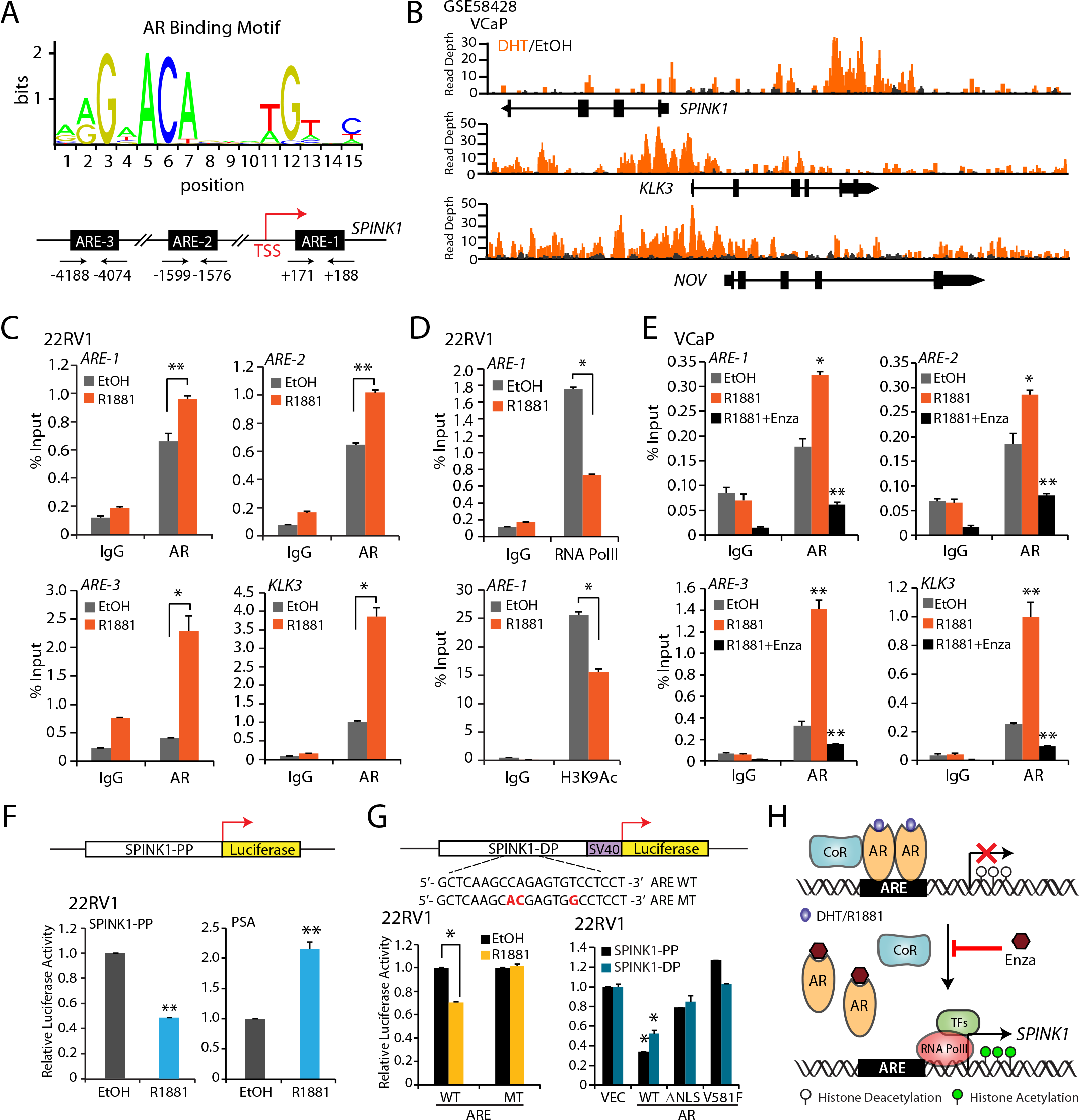
AR directly binds to *SPINK1* promoter and modulates its expression. (A) Schema showing sequence logo for the AR binding motif generated from JASPAR database (top panel). Schema showing genomic location for the Androgen Response Elements (*ARE-1*, *ARE-2* and *ARE-3*) on the *SPINK1* promoter (bottom panel). (B) Chromatin immunoprecipitation (ChIP)-Seq profiles indicating AR enrichment peaks on the *SPINK1*, *KLK3* and *NOV* gene loci in androgen stimulated VCaP cells (GSE58428). (C) ChIP-qPCR data showing recruitment of AR on the *SPINK1* promoter upon androgen R1881 (10nM) stimulation in 22RV1 cells. *KLK3* promoter was used as a positive control for androgen stimulation experiment. (D) ChIP-qPCR data showing occupancy of RNA Pol II on the *SPINK1* promoter in androgen R1881 (10nM) stimulated 22RV1 cells (top panel). ChIP-qPCR data for the H3 lysine 9 acetylation (H3K9Ac) marks under same conditions (bottom panel). (E) ChIP-qPCR data depicting enrichment of AR on the *SPINK1* and *KLK3* promoters in synthetic androgen R1881 (10nM) stimulated VCaP cells treated with or without enzalutamide (10μM). (F) Schema of luciferase reporter construct cloned with proximal promoter of *SPINK1* (SPINK1-PP) upstream of firefly luciferase reporter gene (top panel). Luciferase reporter activity of SPINK1-PP in androgen R1881 stimulated (10nM) 22RV1 cells. PSA promoter construct was used as a positive control for androgen stimulation. (G) Schematic of the luciferase reporter construct cloned with distal promoter of *SPINK1* (SPINK1-DP) wild-type (WT) or mutated (MT) ARE sites (altered residues in red) (top panel). Luciferase reporter activity of SPINK1-DP in androgen R1881 stimulated (10nM) 22RV1 cells (bottom panel, left). Luciferase reporter activity of SPINK1-PP and SPINK1-DP in 22RV1 cells co-transfected with vector control (VEC), AR wildtype (WT) and AR mutants (ΔNLS and V581F) constructs (bottom panel, right). (H) Illustration for AR signaling mediated *SPINK1* regulation in prostate cancer. For all panels except (A), (B) and (I) biologically independent samples were used (n=3); data represents mean ± SEM. \**P*≤ 0.05 and \*\**P*≤ 0.001 using two-tailed unpaired Student’s *t* test.

To confirm AR binding on the *SPINK1* promoter, we performed ChIP-quantitative PCR (ChIP-qPCR) for AR in R1881-stimulated 22RV1 cells. A significant enrichment for AR-binding at three distinct binding sites (*ARE-1*, *ARE-2* and *ARE-3*) was observed in androgen-stimulated 22RV1 cells with respect to EtOH control (Fig 3C and Supplementary Fig S3A). Promoters for *KLK3* and *NOV* were used as controls for the AR binding (Wu et al, 2014). Next, we determined the transcription activity of *SPINK1* by performing ChIP-qPCR for RNA polymerase II (RNA Pol II), interestingly a significant decrease in the occupancy of RNA Pol II on *SPINK1* promoter was noticed in androgen stimulated 22RV1 cells (Fig 3D). Further, a decrease in the recruitment of RNA Pol II on the *NOV* promoter, while no significant change on the *KLK3* promoter was observed upon androgen stimulation in 22RV1 cells (Supplementary Fig S3B). Moreover, ChIP-qPCR for the H3K9Ac, a histone mark for transcriptionally active gene promoters was also performed, and a significant reduction in the enrichment of H3K9Ac activation marks on the *SPINK1* promoter was observed in R1881-stimulated 22RV1 cells, thus confirming its transcriptionally repressed state (Fig 3D). Similarly, a significant increase in the AR-occupancy was observed at all three ARE binding sites on the *SPINK1* promoter in androgen-stimulated VCaP cells, while a remarkable decrease in the androgen-induced AR recruitment was noted in the presence of Enza, indicating impaired AR-binding to these ARE sites in the presence of a potent AR antagonist (Fig 3E). While, no change in the RNA Pol II occupancy on the *SPINK1* promoter was found in androgen stimulated VCaP cells (Supplementary Fig S3D), indicative of its poised transcriptional state.

To further confirm the AR signaling-mediated transcriptional regulation of *SPINK1*, we constructed luciferase promoter reporter vectors by cloning the ARE containing proximal and distal promoter regions of *SPINK1* (SPINK1-PP and SPINK1-DP, respectively) and performed luciferase reporter assay using 22RV1 cells. Upon androgen stimulation, a significant decrease in the luciferase activity was observed in 22RV1 cells transfected with the SPINK1-PP (Fig 3F). While, an increase in the luciferase activity was observed in androgen-stimulated 22RV1 cells transfected with PSA promoter construct, used as a positive control for androgen stimulation. Furthermore, siRNA mediated knockdown of *AR* also led to significant increase in the reporter activity of the SPINK1-PP transfected 22RV1 cells (Supplementary Fig S3E). To identify the critical AR binding site for the transcriptional regulation of *SPINK1*, AREs were mutated (ARE MT) in both SPINK1-PP and SPINK1-DP and luciferase assay was performed. A significant decrease in the luciferase activity was recorded in the 22RV1 cells transfected with ARE WT reporter construct upon androgen stimulation, while mutation of the ARE in the SPINK1-DP showed no change in the luciferase activity (Fig 3G). Conversely, overexpression of wild type AR results in significant decrease in the luciferase activity of both the SPINK1-PP and SPINK1-DP constructs. No significant change in the luciferase activity was observed when AR mutants (AR-ΔNLS and ARV581F) were overexpressed (Fig 3G). In conclusion, our findings revealed that upon androgen stimulation, AR binds to the *SPINK1* promoter and regulates its transcriptional activity by halting the recruitment of RNA Pol II and decreasing the active transcriptional marks (H3K27Ac) on the *SPINK1* promoter. Together these findings support the conclusion that AR functions as a direct transcriptional repressor of the *SPINK1*, and attenuating AR signaling via potent AR-antagonists, such as enzalutamide relieves *SPINK1* transcriptional repression resulting in *SPINK1* upregulation (Fig 3H).

### Increased SPINK1 level promotes epithelial-mesenchymal transition (EMT) and stemness in prostate cancer

To identify the biological processes governed by SPINK1 in PCa cells, we established stable *SPINK1* silenced 22RV1 cells and performed global gene expression profiling. Stable 22RV1 cells transduced with lentivirus-based short hairpin RNAs (shRNAs) (sh*SPINK1*-1, sh*SPINK1*-2 and sh*SPINK1*-3) showed more than ~85% knockdown of *SPINK1* as compared to the control shScramble (shSCRM) cells (Supplementary Fig S4A and B). To further elucidate the functional roles of differentially expressed genes in shSPINK1-1, shSPINK1-2 and shSCRM cells, we performed pathway enrichment analysis using DAVID (Database for Annotation, Visualization and Integrated Discovery). Notably, genes down-regulated upon *SPINK1* knockdown were associated with critical pathways such as, nervous system development, regulation of transcription, and stem cell population maintenance (Fig 4A; Supplementary Table 1). Further, to confirm the role of SPINK1 in EMT, we performed immunostaining for established EMT markers such as E-cadherin (epithelial marker) and Vimentin (mesenchymal marker) in shSCRM and shSPINK1-1 cells. Intriguingly, *SPINK1* silenced cells show a significant increase in the expression of E-cadherin, while a decrease in the expression of Vimentin was observed, indicating that loss of *SPINK1* leads to a decrease in mesenchymal marks in PCa cells (Fig 4B). Previously, we demonstrated the role of SPINK1 in imparting chemoresistance to colorectal cancer cells (Tiwari et al, 2015), thus we investigated whether overexpression of SPINK1 governs the similar attribute in PCa cells. As expected, a significant increase in the chemosensitivity towards well-known chemotherapeutic drugs such as doxorubicin, cisplatin and 5-fluorouracil were recorded in shSPINK1-1 cells as compared to shSCRM cells (Supplementary Fig S4C-E).

**Figure 4.**
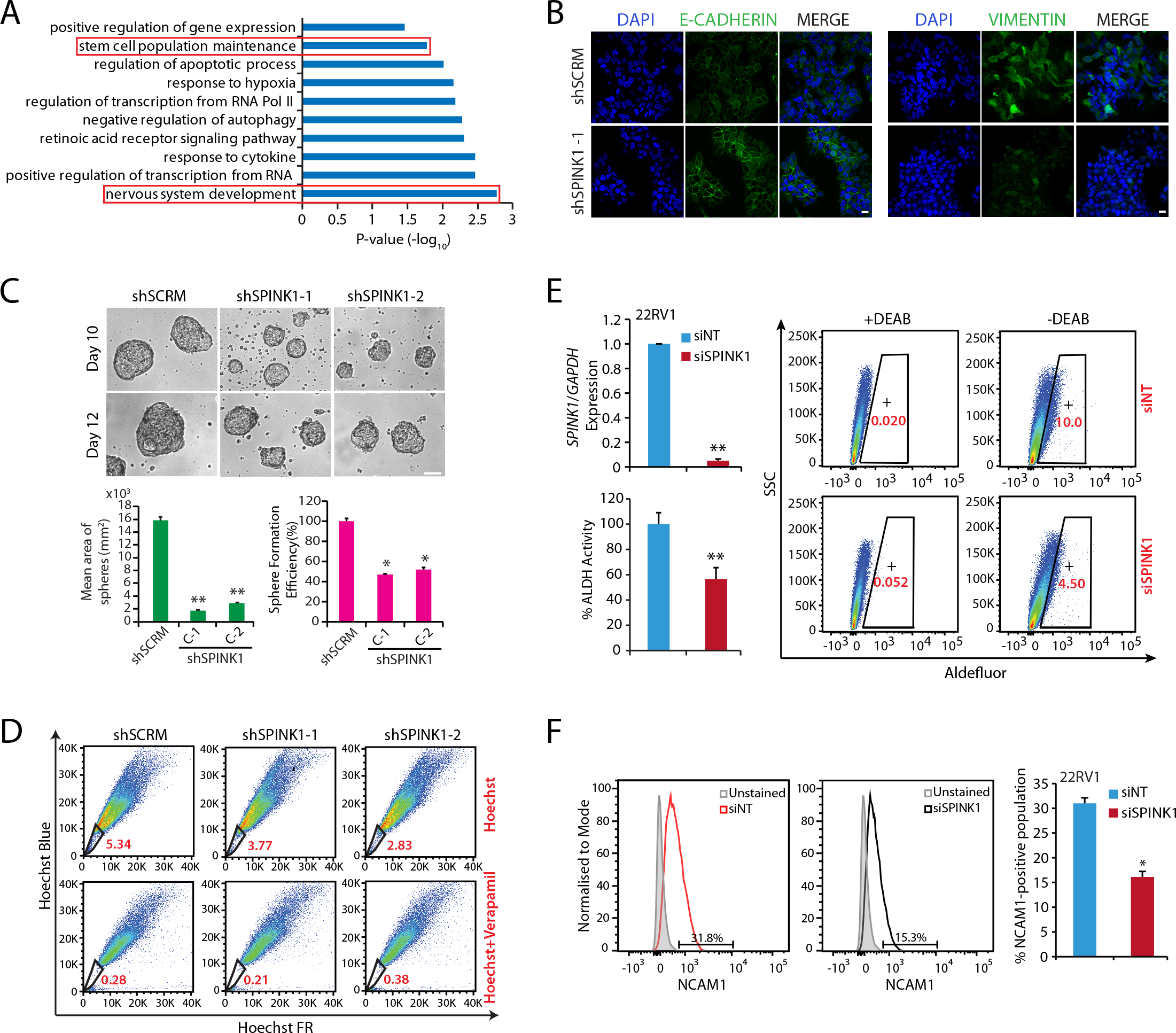
SPINK1 is involved in EMT and maintains chemoresistance in 22RV1 PCa cells. (A) DAVID analysis results showing the gene ontology (GO) associated downregulated pathways in stable *SPINK1* silenced 22RV1 cells relative to control cells (n=3 biologically independent samples). (B) Immunostaining for E-Cadherin (left panel) and Vimentin (right panel) in stable *SPINK1* silenced (shSPINK1-1) and control (shSCRM) 22RV1 cells. Scale bar represents 20 μm. (C) Representative phase contrast microscopic images for the prostatospheres using 22RV1 shSCRM, shSPINK1-1 and shSPINK1-2 cells (top panel). Bar plot depicts percent sphere formation efficiency and mean area of the prostatospheres (bottom panel). Scale bar represents 100 μm. (D) Hoechst-33342 staining for the side population (SP) analysis using same cells as in (C). Percentages of SP were analyzed by putting blue and far red filters, gated regions are marked in red for each panel. (E) ALDH1 expression was measured through flow-cytometry using Aldefluor assay. The top left panel shows quantitative PCR data showing relative expression of *SPINK1* in siRNA mediated *SPINK1* knockdown in 22RV1 cells. The bottom left panel represents the average of five independent quantifications of the ALDH activity in control (si*NT*) and *SPINK1*-silenced 22RV1 cells. The right panel represents flow cytometric graphs showing the fluorescence intensity of catalyzed ALDH substrate in presence and absence of DEAB (ALDH activity inhibitor) in control and *SPINK1* silenced 22RV1 cells. Marked windows in the flow cytometric graphs indicate percentage of ALDH1+ cell population. (F) Flow cytometry histograms depicting NCAM1 expression in control and *SPINK1* silenced 22RV1 cells. The cell populations were normalized to mode. The bar plot represents relative surface expression of NCAM1 in the same cells. In panels (C), (E) and (F) biologically independent samples were used (n=3); data represents mean ± SEM. \**P*≤ 0.05 and \*\**P*≤ 1.1 using two-tailed unpaired Student’s *t* test.

Since, one of the GO terms that showed significant enrichment was maintenance of stem cell population (*P*<0.01) (Fig 4A), thus we performed prostatosphere assay as a readout to test the self-renewal ability of *SPINK1* silenced cells. Noticeably, a significant decrease in the number and size of the prostatospheres was observed in shSPINK1-1, shSPINK1-2 cells as compared to the control cells (Fig 4C). Furthermore, we also performed the side population (SP) assay by evaluating the efflux of Hoechst dye *via* ABC-transporters in the absence or presence of verapamil, a competitive inhibitor for ABC transporters (Zhou et al, 2001). As anticipated, loss of *SPINK1* led to a significant decrease (~29% and ~47%) in the SP in the shSPINK1 cells as compared to control (Fig 4D). Since, aldehyde dehydrogenase (ALDH) activity is crucial for promoting stemness and chemoresistance in cancer stem cells (Burger et al, 2009; Le Magnen et al, 2013), we found a significant decrease in the percent ALDH activity of the *SPINK1*-silenced 22RV1 cells as compared to the control (Fig 4E). Finally, taking a lead from our most significantly enriched GO term underscoring nervous System development (*P*< 0.001) (Fig 4A), we next investigated any alterations in the expression of neuroendocrine prostate cancer (NEPC) markers such as SYP, CHGA, ENO2 in *SPINK1*-silenced 22RV1 cells, notably a significant decrease in the expression of *SYP* (Supplementary Fig S4F) was observed in shSPINK1 cells relative to control, although no change was observed in the CHGA and ENO2 levels (data not shown). Nevertheless, 22RV1 cells with transient *SPINK1* knockdown show a significant reduction in the surface expression of the neural cell adhesion molecule-1 (NCAM1), an established marker of neural lineage and known to induce neurite outgrowth (Fig 4F). Taken together, our findings highlight the predominant role of SPINK1 in mediating EMT, stemness and promoting drug resistance in prostate cancer.

### Androgen deprivation-mediated SPINK1 upregulation is associated with neuroendocrine-like (NE-like) phenotype in prostate cancer

To understand the effect of long-term androgen deprivation on *SPINK1* expression, we investigated publicly available gene expression profiling dataset (GSE8702), wherein LNCaP cells (*SPINK1*-negative) were androgen deprived for 12 months. Remarkably, with prolonged androgen deprivation, a robust increase in the *SPINK1* expression was noticed (Fig 5A). Further, Gene Set Enrichment Analysis (GSEA) revealed that with long-term androgen deprivation, there was a significant decrease in the expression of androgen-signaling associated genes, while positive enrichment of the pathways associated with neuron markers and axon guidance were observed (Supplementary Fig S5A). This further served as an additional piece of evidence emphasizing on the probable role of SPINK1 in cellular plasticity and NE-like morphology. To further confirm the association of SPINK1 overexpression with NE-transdifferentiation, LNCaP cells were cultured in androgen-deprived condition for a duration of 30 days (named as LNCaP-AI cells) (Fig 5B). Consistent with previous report (Yuan et al, 2006), a gradual change in the morphology of LNCaP cells, from an epithelial to a more NE-like phenotype exhibiting neuron-like projections with a concomitant increase in the expression of NEPC markers such as, *SYP*, *CHGA*, *ENO2* and *CD56* was observed (Fig 5B and Supplementary Fig S5B). Furthermore, a significant decrease in PSA and REST levels, while a remarkable increase in the SYP and SOX2 levels was observed in LNCaP-AI cells (Fig 5C). Intriguingly, long-term androgen deprivation also results in a remarkable increase in *SPINK1* expression (~40- and 160-folds up at day 20 and 30, respectively) and corresponding drop in *KLK3* levels in both LNCaP and LNCaP-derived, castration-resistant C4-2 cells (Fig 5C-E and Supplementary Fig S5C). The LNCaP-AI cells also show a significant increase in the expression of EMT (*NCAD*, *VIM* and *TWIST1*) and stemness markers (*CD44* and *KIT*) (Supplementary Fig S5D and E) besides SOX2 (Fig 5C and Supplementary Fig S5E). Next, we also examined for any possible association between *SPINK1* expression and NEPC markers in TCGA-PRAD and MSKCC PCa patient cohorts. Interestingly, a positive correlation between *SPINK1* expression and NEPC markers (*SYP*, *MYCN* and *CHGB*) was observed (Supplementary Fig S5F and G), adding to the evidence on plausible role of SPINK1 in cellular plasticity and reprogramming.

**Figure 5.**
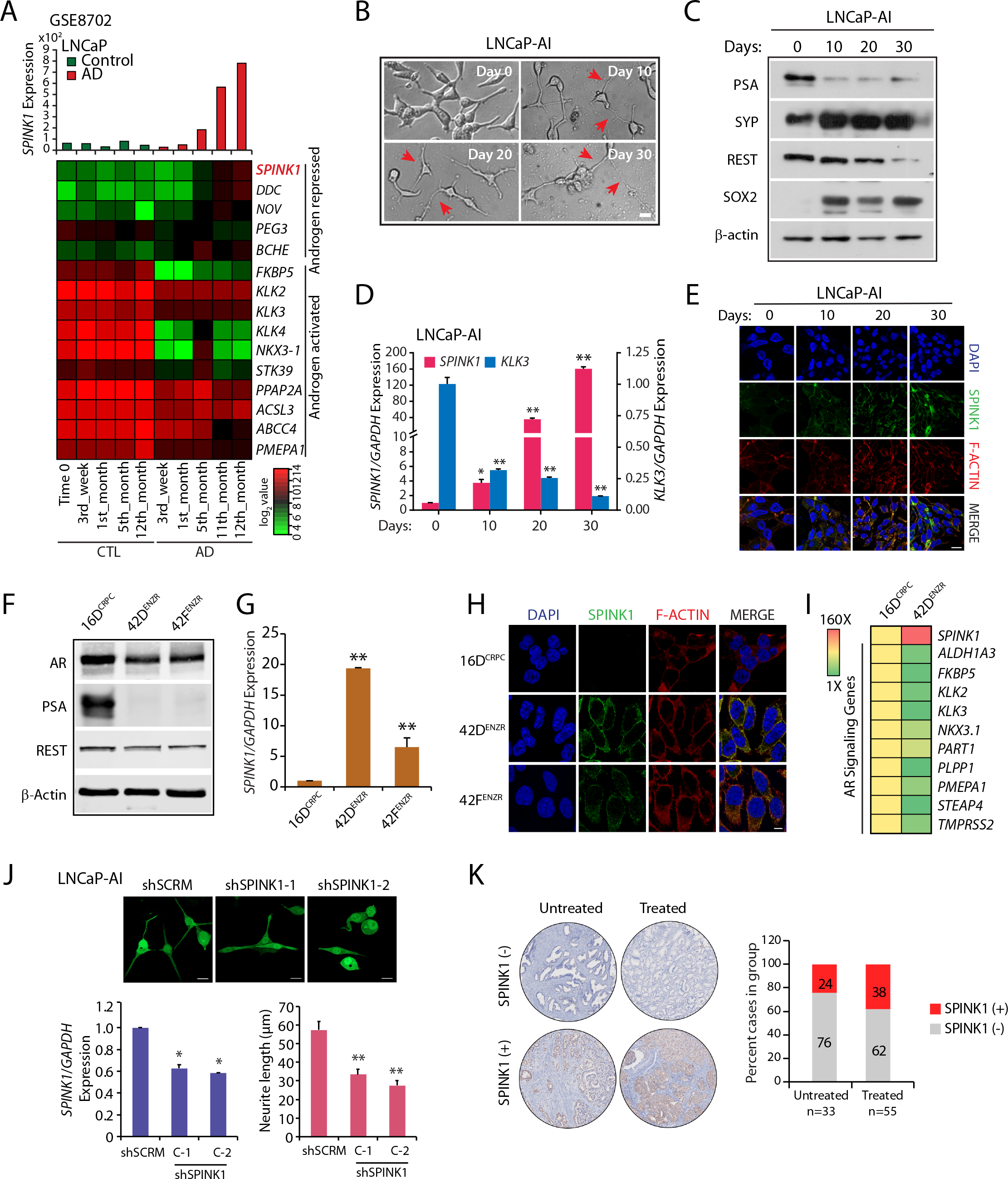
Androgen-deprivation mediated neuroendocrine transdifferentiation of prostate cancer cells show SPINK1 upregulation. (A) Bar graph showing *SPINK1* expression upon long-term androgen deprivation in LNCaP cells (GSE8702). Heatmap depicts expression of AR-signaling associated genes including *SPINK1* in the same LNCaP cells. Shades of red and green represent log2 gene expression values. (B) Representative phase contrast microscopic images for androgen deprived LNCaP cells at indicated time points. (C) Immunoblot assays for PSA, REST, SYP and SOX2 using same cells as in (B). β-actin was included as a loading control. (D) Quantitative PCR data showing relative expression of *SPINK1* and *KLK3* using same cells as in (B). (E) Immunostaining for SPINK1 using same cells as in (B). (F) Immunoblot assay for AR, PSA and REST levels in LNCaP xenografts derivatives, namely 16DCRPC and enzalutamide resistant 42DENZR and 42FENZR cell lines. β-actin was used as a loading control. (G) Quantitative PCR data displaying relative expression of *SPINK1* in the same cells as in (F). (H) Immunostaining for SPINK1 using same LNCaP xenografts derivatives as in (F). Scale bar represents 10μm. (I) Heatmap representing fold increase in *SPINK1* transcript versus AR target genes in 42D^ENZR^ cells compared to 16D^CRPC^. Data is plotted as reads per million. (J) Representative images for the neurite outgrowths in the stable LNCaP shSPINK1 and shSCRM cells, cultured in androgen deprived condition for 30 days (top panel). Bar plot showing quantitative PCR data for relative expression of *SPINK1* (bottom left) and the measurement of neurite lengths in the same cells (bottom right). (K) Representative cores of IHC staining from PCa TMA, comprising of untreated (n=33) and neoadjuvant hormone therapy (NHT) treated patients (n=55). Panel on the top show representative IHC images for SPINK1 expression in SPINK1-negative (SPINK1−) and bottom panel represents SPINK1-positive (SPINK1+) patients. Bar plot showing percentage of cases depicting SPINK1 status in each group. For panels (B), (E) and (J), scale bar represents 20μm. In the panels (D), (G) and (J) biologically independent samples were used (n=3); data represents mean ± SEM. \**P*≤ 0.05 and \*\**P*≤ 0.001 using two-tailed unpaired Student’s *t* test.

To investigate the significance of SPINK1 in governing the cellular plasticity in context of AR signaling, we used LNCaP-derived CRPC cell line, namely 16DCRPC, and its derivative 42D^ENZR^ and 42F^ENZR^ cell lines established via multiple serial transplantation of the enzalutamide resistant tumors in male athymic mice (Bishop et al, 2017). These enzalutamide-resistant cell lines harbor reduced AR activity as depicted by the minimal expression level of PSA as compared to parental 16D^CRPC^ cells (Fig 5F). Further, GSEA plots using the RNA-seq data of 16D^CRPC^ and 42D^ENZR^ cells reveal reduced expression of genes associated with AR signaling, with concomitant increase in the expression of neuronal makers and genes-associated with neurogenesis (Supplementary Fig S5H). Moreover, these enzalutamide resistant cell lines, 42D^ENZR^ and 42F^ENZR^ also show higher expression of NEPC markers such as *SYP*, *CHGA* and *ENO2* relative to the parental line (Supplementary Fig S5I). Notably, 42D^ENZR^ and 42F^ENZR^ cells exhibit increased expression of SPINK1 both at transcript and protein levels as compared to the 16DCRPC cells (Fig 5G and H). Further, transcriptomic analysis revealed negative association of *SPINK1* expression with a panel of classically AR-regulated genes (Bishop et al, 2017; Hieronymus et al, 2006) in 42D^ENZR^ cells, highlighting the inverse association between *SPINK1* expression and AR signaling (Fig 5I). To understand the significance of SPINK1 in these cell lines, we performed siRNA-mediated transient knockdown of *SPINK1* in 42D^ENZR^ and 42F^ENZR^ cells and examined expression of the NEPC markers. Interestingly, loss of *SPINK1* results to a significant decrease in the expression of *SYP* in both the cell lines, while a significant change in the expression of *CHGA* in only 42F^ENZR^ cells was observed (Supplementary Fig S5J and K). No change in the expression of *ENO2* was noted in both the cell lines. To evaluate SPINK1-mediated phenotypic change during NEPC transdifferentiation, we established stable shRNA-mediated *SPINK1* silenced LNCaP cells (LNCaP-shSPINK1) and transdifferentiated these cells in androgen-deprived condition. Intriguingly, LNCaP-shSPINK1 showed significant reduction in the length of neurite-like projections as compared to control SCRM cells (Fig 5J), indicating the significance of SPINK1 in NEPC transdifferentiation and cellular plasticity.

To further examine the effect of ADT on the SPINK1 expression in clinical samples, we examined the expression of SPINK1 in a TMA comprising of PCa patient specimens (n=88) by performing IHC staining, wherein 55 out of 88 patients were given neoadjuvant hormone therapy (NHT) for 3 months. In concordance with our in vitro findings including enzalutamide-resistant tumors derived cells, about ~38% (21 out of 55) patients who were administered NHT exhibits SPINK1 positive status compared to only ~24% (8 out of 33) in the untreated group (Fig 5K). Although, ADT or NHT-mediated SPINK1 upregulation and associated risk factors need to be tested in a larger PCa patient cohort. Collectively, our finding shows a trend that androgen-deprivation therapies may have an adverse effect, and the benefits must be weighed against treatment. Conclusively, we also show that elevated SPINK1 levels during NE-transdifferentiation strongly emphasizes the potential role of SPINK1 in governing stemness and cellular plasticity in prostate cancer.

### SPINK1 expression is modulated by reprogramming factor SOX2 and AR transcriptional co-repressor REST

The role of SRY (sex determining region Y)-box 2 (SOX2) has been implicated in neuroendocrine differentiation and reprogramming/lineage plasticity in *RB1* and *TP53* deficient prostate cancer (Mu et al, 2017; Russo et al, 2016). Moreover, SOX2 has also been known as an androgen repressed gene (Kregel et al, 2013). Since, our data also showed a similar trend of decrease in SOX2 expression as SPINK1 in androgen-stimulated 22RV1 cells (Supplementary Fig S6A), we sought to examine SOX2 mediated regulation of *SPINK1* expression. We scanned the *SPINK1* promoter for the SOX2 binding motif using MatInspector (Cartharius et al, 2005), and identified three putative binding sites (*S1*, *S2* and *S3*) adjacent to the TSS (Fig 6A). To confirm SOX2 binding on the *SPINK1* promoter, we performed ChIP-qPCR in 22RV1 cells, an endogenous SOX2 positive cell line, interestingly a significant enrichment for SOX2-binding at all the three distinct binding sites was observed (*S1*, *S2* and *S3*) (Fig 6B). To ascertain the transcriptional significance of SOX2 binding, we also looked for RNA Pol II binding on these sites and found an enrichment in the occupancy of RNA Pol II on the *SPINK1* promoter (Fig 6C). However, no change in *SPINK1* expression was observed upon siRNA-mediated knockdown of *SOX2*, suggesting that other regulatory factor(s) might be involved in governing *SPINK1* expression (Supplementary Fig S6B). To further investigate whether increase in the SPINK1 level in LNCaP cells during NE-like transdifferentiation is transcriptionally regulated by SOX2, we examined for SOX2 recruitment on the *SPINK1* promoter region using LNCaP cells cultured in androgen deprived condition (LNCaP-AI) for 15 days. Interestingly, we found a remarkable enrichment of SOX2 on the *SPINK1* promoter in LNCaP-AI cells as compared to the normal LNCaP cells grown in regular media (Fig 6D). In addition, an increase in the occupancy of RNA Pol II was also noticed on the *SPINK1* promoter in LNCaP-AI cells (Fig 6E), signifying the increased transcriptional activity of *SPINK1* promoter. Next, to further confirm that SOX2 positively regulates SPINK1, we overexpressed SOX2 in LNCaP cells (SPINK1-negative), and examined for the expression of SPINK1, in agreement, a significant increase in the *SPINK1* expression was observed in SOX2 overexpressing LNCaP cells relative to the control cells (Fig 6F). Moreover, a significant increase in the luciferase activity of SPINK1-DP promoter was observed in the SOX2 overexpressing LNCaP cells (Fig 6G).

**Figure 6.**
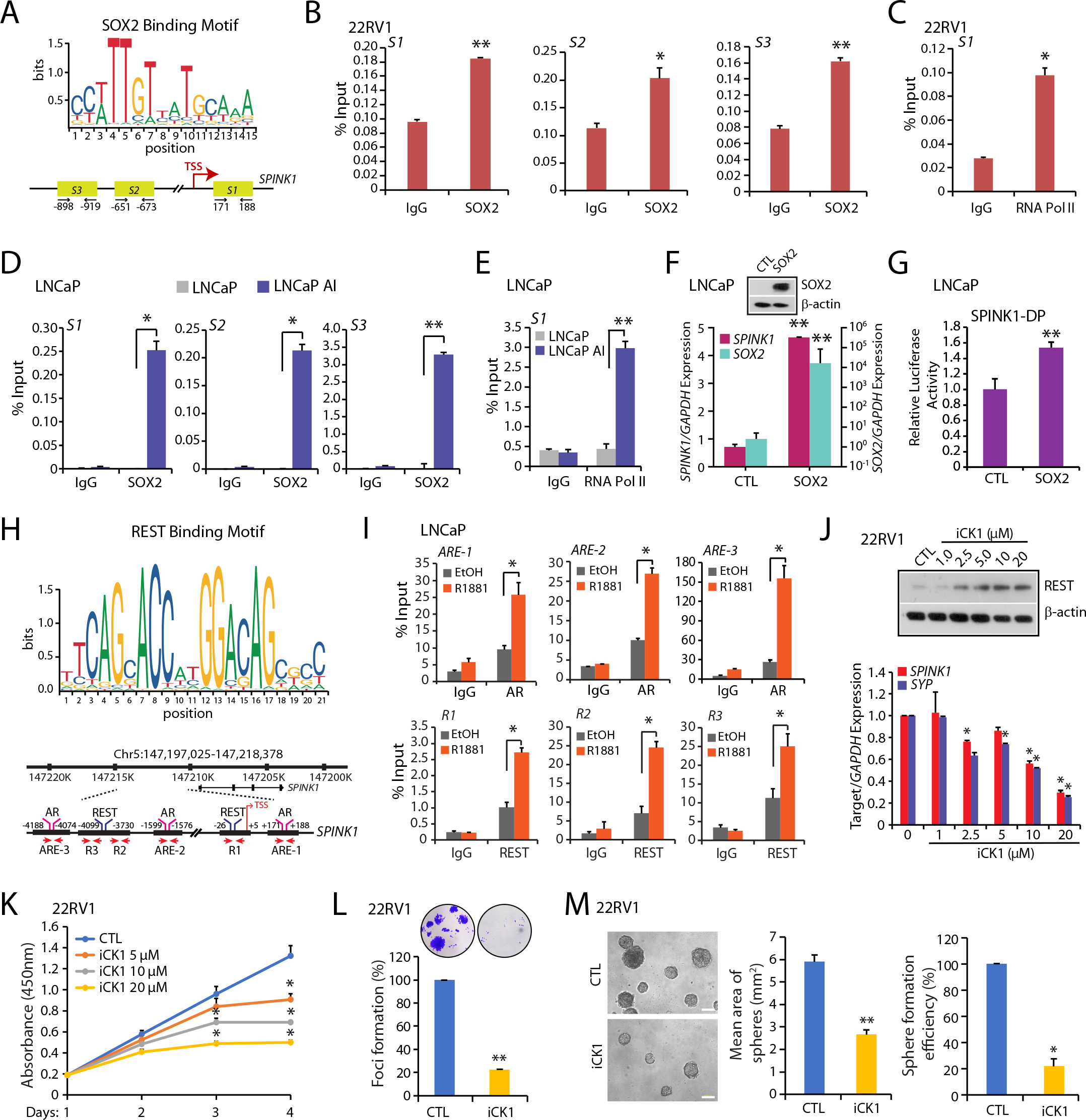
Reprogramming factor SOX2 and AR transcriptional co-repressor REST modulate *SPINK1* expression. (A) Schema showing the sequence logo for the SOX2 binding motif constructed from JASPAR database (top panel). Schematic positioning of SOX2 binding elements (*S1*, *S2* and *S3*) on the *SPINK1* promoter region (bottom panel). (B) ChIP-qPCR data showing occupancy of SOX2 on the *SPINK1* promoter in 22RV1 cells. (C) Same as in (B), except recruitment of RNA Pol II is shown. (D) ChIP-qPCR data showing occupancy of SOX2 on the *SPINK1* promoter in normal LNCaP and LNCaP-AI cells (androgen deprived for the period of 15 days). (E) ChIP-qPCR data for RNA Pol II binding on the *SPINK1* promoter using the same cells as in (D). (F) Immunoblot assay for in SOX2-overexpressing LNCaP cells (top panel). Quantitative PCR data showing relative expression of *SOX2* and *SPINK1* upon transient SOX2 overexpression in LNCaP cells (bottom panel). β-actin was used as a loading control. (G) Luciferase reporter activity of the SPINK1 distal-promoter (SPINK1-DP) in LNCaP cells transfected with vector control (CTL) or SOX2 overexpression constructs. (H) Schema showing the sequence logo for the REST binding motif constructed from JASPAR database (top panel). Genomic location for AR and REST binding sites on the *SPINK1* promoter region and positioning of the ChIP-qPCR primers (bottom panel). (I) ChIP-qPCR data showing enrichment of AR and REST on the *SPINK1* promoter in androgen R1881 stimulated (10nM) LNCaP cells. (J) Immunoblot analysis for the REST level in 22RV1 cells treated with Casein Kinase 1 inhibitor (iCK1) at the indicated concentrations (top panel). β-actin was used as a loading control. Quantitative PCR data for relative *SPINK1* and *SYP* expression for the same cells (bottom panel). (K) Cell proliferation assay using 22RV1 cells treated with different concentrations of iCK1 at the indicated time points. (L) Foci formation assay using 22RV1 cells treated with iCK1, 20μM concentration. Representative images depicting foci are shown in the inset. (M) Representative phase contrast microscopic images of 3D tumor spheroid assay using the same cells as in (K) are shown at the left. Bar plots depict mean area of spheres and sphere formation efficiency. Scale bar represents 1000μm. In all panels except (A) and (H), biologically independent samples were used (n=3); data represents mean ± SEM. \**P*≤ 0.05 and \*\**P*≤ 0.001 using two-tailed unpaired Student’s *t* test.

Downregulation of transcriptional repressor REST has been well-established phenomenon involved in the transdifferentiation of CRPC to NEPC phenotype (Lapuk et al, 2012). On the other hand, REST is also known to act as a transcriptional co-repressor for AR for a subset of genes involved in promoting NE-like phenotype (Svensson et al, 2014). Since, we have already established that the AR signaling plays critical role in transcriptional repression of *SPINK1*, and upregulation of SPINK1 positively correlates with NE-like phenotype, thus we next examined the plausible association of SPINK1 with REST and with other members of REST complex using TCGA-PRAD dataset. Quartile-based normalization method was used to stratify the patients based on high and low *SPINK1* expression, notably SPINK1-high patients (SPINK1-positive) show inverse correlation between expression of *SPINK1* and *REST* as well as other members of REST complex such as *RCOR1*, *SIN3A*, *HDAC1* (Supplementary Fig S6C and D). Next, we sought to examine the role of REST in AR-mediated transcriptional repression of SPINK1 in PCa cell lines. Notably, androgen stimulation in three different PCa cell lines, 22RV1, LNCaP and VCaP results in a significant increase in the REST expression (Supplementary Fig S6E), while abrogating the AR signaling using AR-antagonists in VCaP cells resulted in significant decrease in REST expression (Supplementary Fig S6F). To investigate whether REST is acting as a transcriptional co-repressor of AR in the regulation of *SPINK1*, we examined *SPINK1* promoter for the REST binding motif using MatInspector (Cartharius et al, 2005), and checked for the recruitment of both AR and REST within ~5Kb region of the TSS of the *SPINK1* gene (Fig 6H). As expected, a robust enrichment of AR at all the three distinct ARE binding sites (*ARE-1*, *ARE-2* and *ARE-3*) was observed in androgen-stimulated LNCaP cells with respect to EtOH (Fig 6I); intriguingly, a remarkable recruitment of the REST was also observed at the three distinct RE1 sites (*R1*, *R2* and *R3*) adjacent to the AR occupied ARE sites (Fig 6I and Supplementary Fig S6G). Moreover, REST is post-translationally regulated by F-box protein E3 ubiquitin ligase SCF (β-TrCP) responsible for its degradation (Kaneko et al, 2014; Westbrook et al, 2008). Phosphorylation of serine residues of the non-canonical degron motifs in the C-terminal of REST by Casein Kinase 1 (CK1) enables its binding to the E3 ubiquitin ligase β-TrCP resulting in ubiquitin-mediated proteasomal degradation of REST in hippocampal neurons (Kaneko et al, 2014). Thus, to confirm the role of REST as the transcriptional corepressor of AR, we treated 22RV1 (SPINK1+) cells which exhibit low REST expression (Chang et al, 2017b) with a CK1 inhibitor (iCK1, D4476), notably a significant increase in REST levels was observed, subsequently resulting in more than ~70% decrease in the *SPINK1* expression at the highest concentration of iCK1 (Fig 6J), along with a concomitant decrease in *SYP* expression (Fig 6J). To examine the functional relevance of CK1 inhibition, we treated 22RV1 cells with a range of iCK1 concentrations, and a significant reduction in the cell viability was observed at higher concentration (10μM and 20μM) of iCK1 (Fig 6K). Similarly, iCK1 treatment also diminished the foci forming ability of the 22RV1 cells (Fig 6L). To evaluate its effect on the neoplastic transformation ability of 22RV1 (SPINK1+) cells, we also performed three-dimensional tumor spheroid assay, intriguingly a significant reduction in the number and size of the tumor spheroids was observed with the highest concentration of iCK1 (Fig 6M). In summary, our findings suggest that the iCK1 stabilizes the REST levels, which in cooperation with AR, elicits transcriptional repression of SPINK1 and inhibits SPINK1-mediated oncogenic properties.

Collectively, we have shown the direct role of SOX2 in the transcriptional regulation of *SPINK1* in prostate cancer. We also establish that REST acts as a transcriptional corepressor of AR in modulating *SPINK1* expression, thus a cease in the AR signaling during NE-transdifferentiation results in SPINK1 upregulation, and its overexpression positively associates with NE-like phenotype (Fig 7).

**Figure 7.**
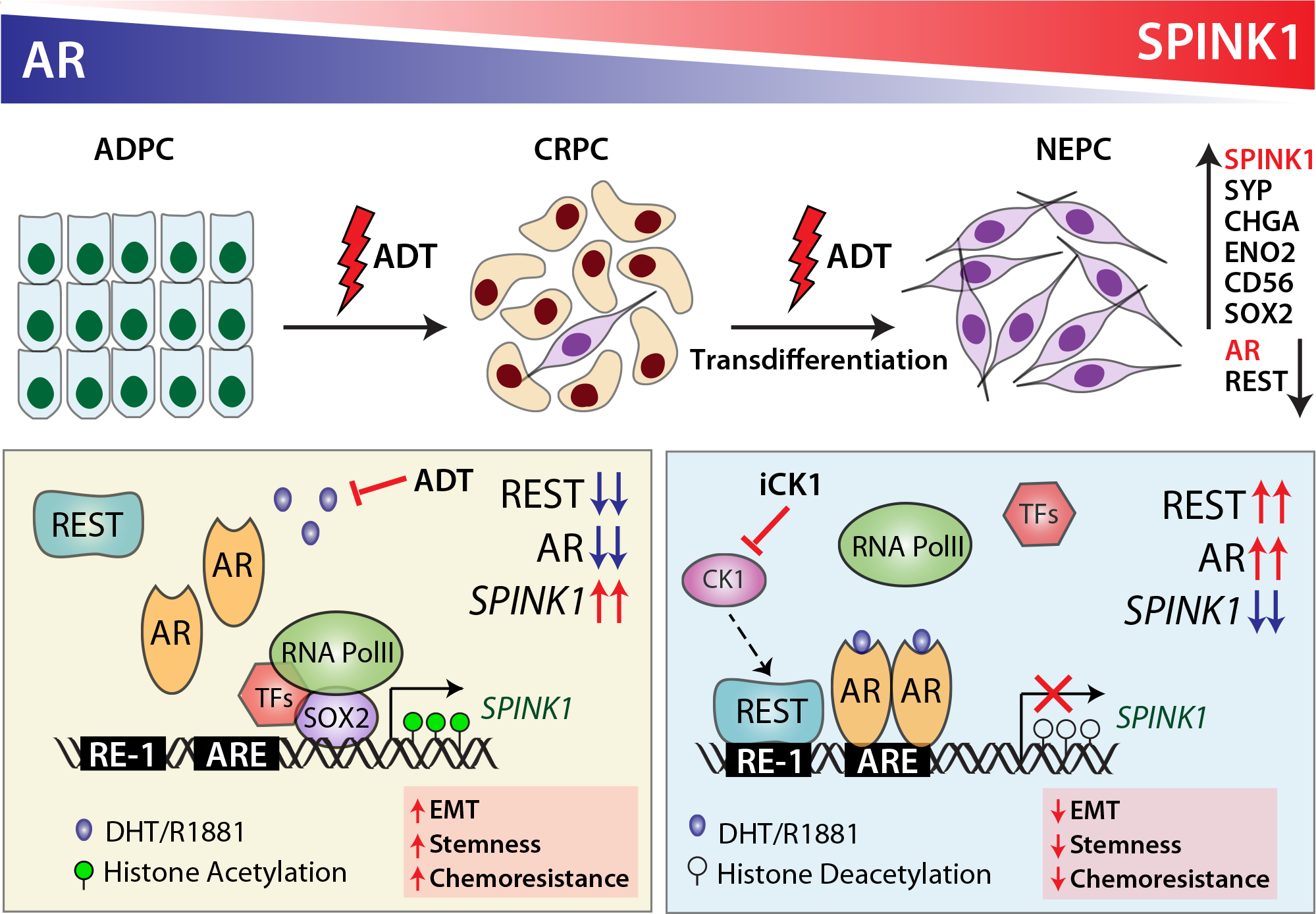
Androgen receptor and its transcriptional co-repressor REST modulate *SPINK1* expression. Schematic representation of AR-REST mediated transcriptional repression governing the expression of *SPINK1*. Androgen deprivation therapy (ADT) using potent AR-antagonists relieve this repression resulting in *SPINK1* upregulation. Casein kinase 1 inhibitor enhances REST levels leading to reduced *SPINK1* expression, resulting in the reduced stemness and cellular plasticity.

## Discussion

Overexpression of SPINK1 in prostate cancer has been associated with poor prognosis including decreased response to ADT, faster progression to castrate-resistant stage and cancer-associated mortalities, thus underpinning its significance as an over-all biomarker of aggressivity and an indicator for poor clinical response. A recent study showed that exogenous expression of HNF4G or HNF1A activates gastrointestinal-lineage transcriptome in PCa, and results in the upregulation of numerous PCa-gastrointestinal signature genes including SPINK1 (Shukla et al, 2017). However, the exact mechanism how SPINK1 is regulated in prostate cancer, and why its overexpression is often associated with aggressive phenotype remains unclear. In this study, we provide a compelling evidence that *SPINK1* is an androgen-repressed gene, and androgen deprivation therapies, using AR antagonists relieve the AR signaling-mediated repression of *SPINK1* resulting in its overexpression. The prominent recruitment of REST at three distinct RE1 sites adjacent to the AR occupied ARE sites on the *SPINK1* promoter in androgen-stimulated LNCaP cells, confirms its role as an AR transcriptional co-repressor involved in the regulation of *SPINK1*. Furthermore, immunostaining of PCa specimens further confirmed an inverse association between AR and SPINK1 expression. A recent study showed that anatomic location of the tumor in the prostate gland is influenced by AR signaling, for instance anterior tumors tend to have lower global AR signaling leading to differences in AR molecular sub-types, tumor size, and PSA (Faisal et al, 2016). Interestingly, African-American men with aggressive PCa largely of SPINK1-positive subtype show higher propensity for anteriorly localized tumors as compared to Caucasian men with matched clinicopathologic features (Faisal et al, 2016; Sundi et al, 2014). Previously, Paju et al first time demonstrated reduced secretion of TATI (SPINK1) in 22RV1 cells upon androgen stimulation (Paju et al, 2007). They also showed elevated SPINK1 level to be associated with higher Gleason grade, and higher expression of neuroendocrine marker, CHGA in a subset of cells (Paju et al, 2007). Taken together, these independent reports further strengthen our findings that SPINK1 is an androgen-repressed gene, which show negative association with AR and other members of the AR repressor complex. Beltran and colleagues showed distinct concordance with respect to the *TMPRSS2-ERG* fusion status in NEPC vs. PCa specimens, wherein NEPC foci failed to show ERG protein expression even in the tumors harboring *TMPRSS2-ERG* rearrangement, directing towards the absence of AR expression, and further confirming that expression of ERG oncoprotein from *TMPRSS2-ERG* fusion transcript is controlled by androgen signaling (Beltran et al, 2011). Since, multifocality remains a matter of concern in PCa molecular subtyping, such that SPINK1 expression is found to be restricted to a few or more foci (Smith & Tomlins, 2014), thus we speculate that low AR signaling in some foci (such as those within anterior lobes) of prostate gland may results in SPINK1 overexpression (SPINK1-positive focus), which by acquiring critical alterations (overexpression of *AURKA* and *MYCN*) drive the neuroendocrine transdifferentiation.

Androgen receptor is a well-defined transcriptional activator for numerous oncogenes involved in the neoplastic transformation of prostatic cells. However, apart from its conventional tumor promoting activity, it has also been identified as a tumor suppressor (Gao et al, 2016). Androgen regulated genes, such as *KLK3*, *KLK2*, *NKX3A*, *TMEPA1*, *TMPRSS2* and *SPAK* represents important features of AR signaling in prostate, while same set of genes also play critical role in diverse physiological processes in the neoplastic progression (Nelson et al, 2002). Hence, targeting androgen signaling by surgical or chemical castration remains the primary therapeutic modality for the treatment of advanced stage PCa patients. Further, the mechanistic insights into androgen signaling led to the development of several novel therapeutic interventions, often with limited success for men with castrate-resistant disease (Claessens et al, 2014). Recently, it has been reported that E3 ubiquitin ligase, SIAH2 regulate ubiquitination-dependent degradation of multiple substrates, including nuclear corepressor AR/NCOR1 complex, and regulates the expression of a subset of AR target genes during CRPC development (Lopez et al, 2016; Qi et al, 2013). Moreover, homozygous deletion of *Siah2* in the dorsal prostate of *TRAMP* mice results in decreased *Spink3* (a mouse homologue of *SPINK1*) levels along with other AR regulated genes including *NKX3.1 (Qi et al, 2013)*. The REST, also known as Neuron-Restrictive Silencer Factor (NRSF) belongs to Kruppel-type zinc finger transcription factor family (Tapia-Ramírez et al, 1997), acts as a master negative regulator of the genes involved in neuronal differentiation (Gao et al, 2011). Moreover, REST is also known to mediate gene repression by recruiting CoREST and SIN3A by binding to the RE1 elements of the target genes, which in turn recruits histone deacetylases, HDAC-1/-2, thereby governing epigenetic reprogramming (Noh et al, 2012). Therefore, it is likely that AR and REST repressive complex may also involve other co-factors such as CoREST in repressing the *SPINK1* expression. In another recent study, REST is shown to be a downstream effector of PI3K/AKT signaling, and inhibitors targeting this signaling axis results in enhanced degradation of REST through-TRCP mediated β proteasome pathway, which in turn promote NE transdifferentiation of PCa cells (Chen et al, 2017). Further, FOXA1 is known to recruit the LSD1-CoREST complex to androgen-regulated enhancers, wherein this complex function through HDAC-1 and −2, and suppress basal transcription of the target genes even in the absence of AR (Cai et al, 2014). Inactivation of the transcriptional repressor REST in PCa cells show upregulation in the expression of neuronal specific genes namely, *CHGA* and *SYP* (Svensson et al, 2014). Furthermore, REST also suppresses EMT and stemness by repressing the expression of genes, namely CD44 and TWIST1 in 22RV1 cell line which is known to exhibit NE-like phenotype (Chang et al, 2017a). In the present study, we have identified REST to be a transcriptional corepressor of AR, which negatively regulates SPINK1. Moreover, we also found inverse association between *SPINK1* and *REST* as well as other members of the REST interacting complex such as *RCOR1*, *SIN3A* and *HDAC1* in PCa patients (Supplementary Fig S6C). Importantly, we show that targeting the ubiquitination-dependent REST degradation using Casein Kinase 1 inhibitor (iCK1) results in reduced *SPINK1* and *SYP* expression, subsequently exhibiting decrease in oncogenic properties. Collectively, our finding suggests that AR along with REST, a transcriptional co-repressor modulate the expression of *SPINK1*, thus stabilizing the REST protein levels suggest a novel therapeutic strategy in controlling the SPINK1-mediated oncogenicity.

Ablation of androgen signaling has been implicated in upregulation of several EMT markers (*SNAI1*, *SNAI2*, *CDH2*, *ZEB1* and *TWIST1*), a phenotype often associated with PCa metastases (Jennbacken et al, 2010; Sun et al, 2012a; Zhifang et al, 2015; Zhu & Kyprianou, 2010). Further, a bidirectional negative feedback loop between AR and ZEB1 has been established, which drives EMT and stem cell–like features following androgen deprivation in LuCaP35 prostate tumor explants (Sun et al, 2012a). Similarly, androgen deprivation in *PTEN* deficient PCa mouse model show faster progression of high-grade PIN (HGPIN) to an invasive phenotype (Jia et al, 2013). In the present study, we show that *SPINK1* knockdown in 22RV1 cells exhibit decrease in EMT, stemness and enhanced chemosensitivity, which could be attributed to decrease in the side-population as well as ALDH activity (Fig 4E), thus indicating the role of SPINK1 in imparting chemoresistance, stemness and cellular plasticity. Recently, we have shown that SPINK1 expression positively correlates with EZH2, a member of Polycomb repressive complex 2, known to induce pluripotency and stemness (Bhatia et al, 2018). Furthermore, SOX2 has been implicated as a key regulator in governing pluripotency, neural differentiation (Zhang & Cui, 2014), and being an AR repressed gene, it is also known to drives NE-transdifferentiation (Mu et al, 2017). Notably, knockdown of SOX2 in mouse embryonic cells result in down-regulation of *Spink3* (Sharov et al, 2008). In our study, we for the first time demonstrate that overexpression of SOX2 in LNCaP cells (SPINK1-negative) results in upregulation of SPINK1, indicating SOX2-mediated positive regulation of *SPINK1*. Further, a significant recruitment of SOX2 on the *SPINK1* promoter leading to upregulation of SPINK1 in 22RV1 as well as androgen deprived LNCaP cells further confirms SOX2-mediated transcriptional regulation of *SPINK1*. Collectively, we propose a novel mechanism involved in SPINK1 regulation, wherein decreased AR and REST levels relieve the repression of the *SPINK1* promoter, subsequently SOX2 gets recruited onto *SPINK1* promoter resulting in its enhanced transcriptional activity.

Conclusively, our findings emphasize that administering PCa patients with androgen deprivation therapies, may result in increased SPINK1 levels accompanied by upregulation of NE markers, thus escalating ADT-associated NEPC incidence (Fig 7). Although, androgen ablation therapy is well-established for the treatment of PCa patients, its long-term benefits are still debatable, but several studies have identified EMT and metastasis as potential disregarded consequences of the potent AR-antagonists used in the clinics. Thus, there is an obligation to take a well-informed decision and the ADT benefits must be weighed before administering anti-androgens, or drug-regimens involving AR-antagonists.

## Materials and Methods

### Human Prostate Cancer Specimens

Tissue microarrays (TMA) with prostate cancer (PCa) specimens were obtained from the Department of Pathology, Henry Ford Health System, Detroit, Michigan, USA and Vancouver Prostate Centre (VPC), Vancouver, BC, Canada after acquiring the due consent from the patients and mandatory approval from the Institutional Review Board. All patients’ specimens used in this study were collected in accordance to the Declaration of Helsinki. The TMA comprises of cases with radical prostatectomy, mostly with localized cancer and some may have lymph node metastases. The VPC TMA comprises of hormone naive cases (n=33) and cases administered with neoadjuvant hormone therapy (n=55), comprising of LHRH agonists and bicalutamide for 3 months. The TMAs were stained for SPINK1 and AR using immunohistochemistry (IHC).

### Evaluation criteria for SPINK1 and AR expression

The IHC staining for SPINK1 was evaluated as positive or negative as described previously (Bhalla et al, 2013). The staining for AR was scored into four different categories: high, medium, low and negative, based upon staining intensity. Further, association between SPINK1 and AR expression in patients’ samples was inferred by applying Chi-square test and Fisher's exact test on GraphPad Prism.

### AR signaling score

AR signaling score was obtained using previously published AR gene signature (Bluemn et al, 2017; Hieronymus et al, 2006). Firstly, PCa patients from TCGA-PRAD and MSKCC cohorts were stratified by performing quartile-based normalization to classify the patients based on high and low *SPINK1* expression. For the signaling score, gene expression of each gene mentioned in the AR gene signature was downloaded as a respective Z-score from cBioPortal and the signaling score was then determined by computing the mean expression of these genes. For both the cohorts, signaling score between the top and bottom quartile was compared. Significance of the signaling score among *SPINK1*-high and low patients was evaluated by unpaired, two-tailed Student’s *t*-test on GraphPad Prism.

### Statistics

Statistical significance was determined by unpaired, two-tailed Student’s *t*-test for independent samples or otherwise mentioned. The differences between the different groups were considered significant if the *P*-value was less than 0.05. Significance is indicated as follows: \**P*<0.05, \*\**P*<0.001. Error bars represent standard error of the mean (SEM) obtained from experiments performed at least three independent times.

### Data availability

The gene expression microarray data from this study has been submitted to the NCBI Gene Expression Omnibus (GEO, http://www.ncbi.nlm.nih.gov/geo/) under the accession number GSE124345.

## Acknowledgements

B.A. is an Intermediate Fellow of the Wellcome Trust/DBT India Alliance. This work is supported by the Wellcome Trust/DBT India Alliance Fellowship [grant number: IA/I(S)/12/2/500635] awarded to B.A. R.T. thanks the Indo-Canadian Shastri Research Student Fellowship (SRSF) for short-term travel support. We are grateful to Prof. Noel Buckley for his insightful suggestions. We thank Anjali Bajpai for critically reading the manuscript.

## Authors’ contributions

R.T., N.M., and B.A. designed and directed the experimental studies. R.T. and N.M. performed *in-vitro* cell line-based studies. R.T., N.M. and V.B. performed the gene expression studies, bioinformatics analysis and ChIP assays. R.T. and A.Y. performed the immunofluorescence. R.T., N.M. and B.A. performed statistical analysis and interpreted the data. A.Z provided the neoadjuvant hormone therapy (NHT) tissue microarray data, LNCaP xenograft-derived cell lines and helped in designing ALDH and NCAM1 experiments. S.C., N.G., and N.P. assembled PCa tissue microarrays and performed immunohistochemistry staining. R.T., N.M. and B.A. wrote the manuscript. B.A. directed the overall project.

