## Supplementary Figures and Methods for "Androgen deprivation upregulates SPINK1 expression and potentiates cellular plasticity in prostate cancer"

<u>**Supplemental Data**</u>

**Financial support:** This work is supported by the Wellcome Trust/ DBT India Alliance grant (IA/I(S)/12/2/500635 to BA).

**Conflict of interest:** The authors declare no conflicts of interest or disclosures.

***Corresponding Author**:
Bushra Ateeq, Ph.D.
Molecular Oncology Lab,
Department of Biological Sciences and Bioengineering,
Indian Institute of Technology Kanpur, Kanpur, 208016, U.P., INDIA
 (B.A.)

### Supplementary Figure S1

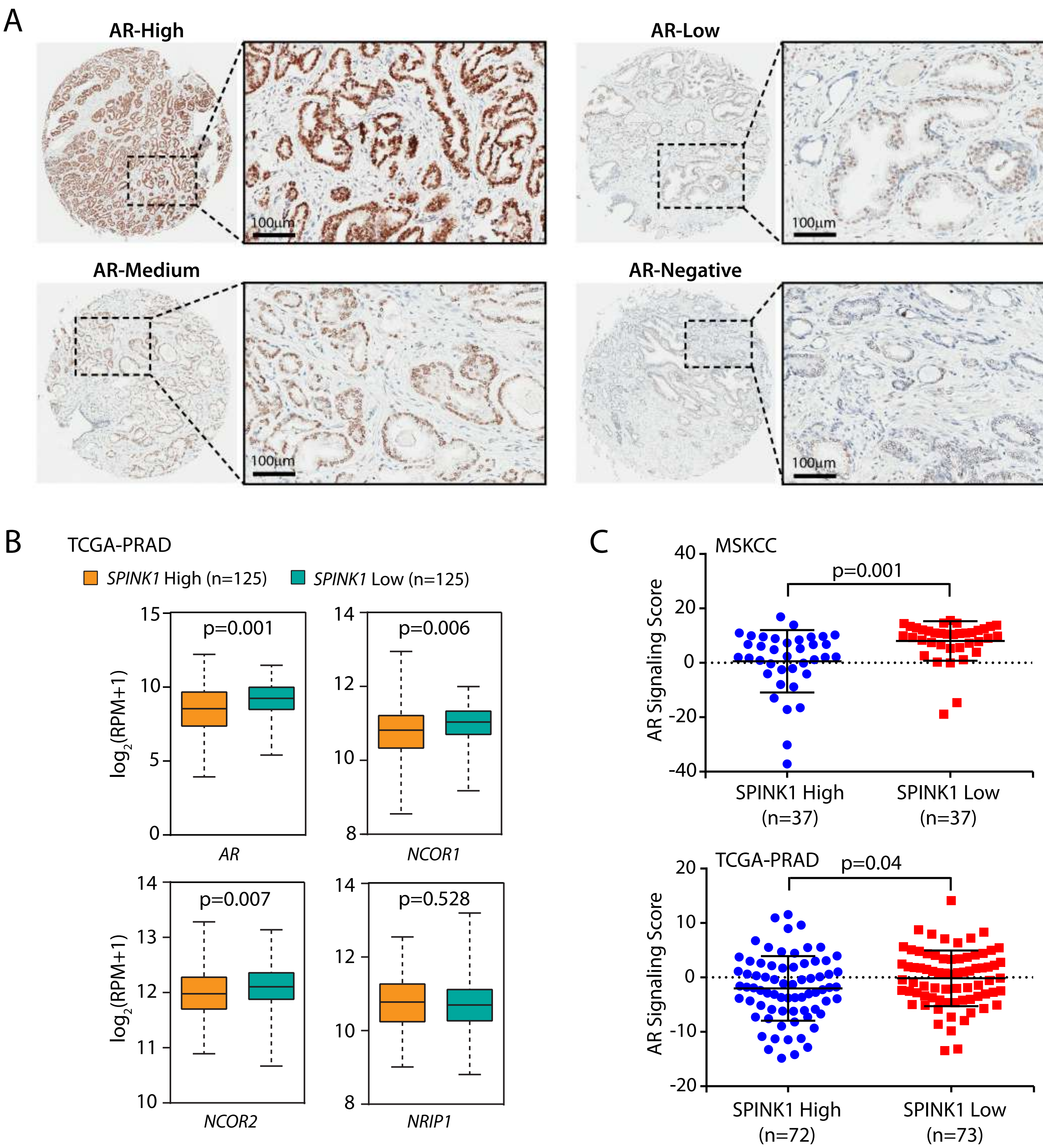


**Supplementary Figure S1. *SPINK1* expression is negatively correlated with *AR* and members of AR repressive complex in PCa patients.**

**(A)** Representative micrographs showing immunohistochemical (IHC) staining for AR in PCa tissue microarray (TMA) cores (n=237), wherein based on intensity AR staining is categorized as high, medium, low and negative (Gleason score 7). Scale bar represents 500 µm and 100 µm for the entire core and the inset, respectively. **(B)** Relative expression of the members of AR repressive complex (*AR*, *NCOR1*, *NCOR2* and *NRIP1*) in SPINK1 high (n=125) and low (n=125) PCa patients, stratified by employing quartile based normalization of SPINK1 expression in TCGA-PRAD dataset. Transcripts level shown as $\log_2$ (RPM+1). **(C)** Dot plot depicting inverse association between AR signaling score and *SPINK1* expression in the MSKCC (n=74; top) and TCGA-PRAD datasets (n=145; bottom) in SPINK1 high and low PCa patients, stratified by employing quartile based normalization of SPINK1 expression. AR signaling score represents the mean expression of 10-gene signatures linked to AR signaling.

In panels (B) and (C), *P*-values were calculated using two-tailed unpaired Student's *t* test.

### Supplementary Figure S2

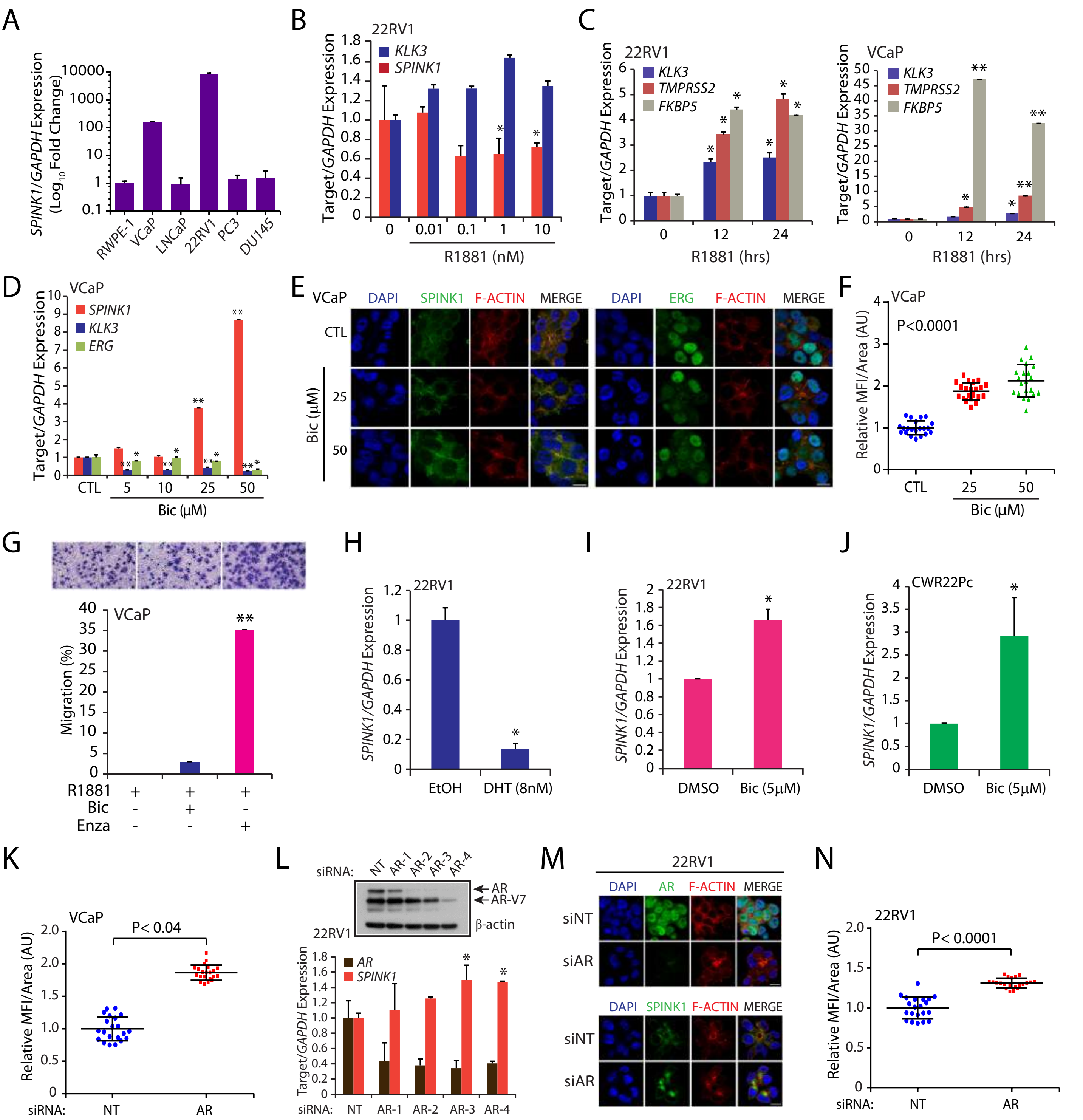


**Supplementary Figure S2. SPINK1 expression is negatively regulated by androgen signaling in PCa cells.**
**(A)** Quantitative PCR data showing relative expression of *SPINK1* in the PCa cell lines panel. **(B)** Quantitative PCR data showing relative expression of *SPINK1* and *KLK3* in 22RV1 cells stimulated with androgen (R1881) at various concentrations. **(C)** Quantitative PCR data showing relative expression of androgen regulated genes (*KLK3*, *TMPRSS2* and *FKBP5*) in androgen (R1881, 10 nM) stimulated 22RV1 and VCaP cells at different time points. **(D)** Quantitative PCR data showing relative expression of *SPINK1*, *KLK3* and *ERG* in bicalutamide (5, 10, 25 and 50μM) treated VCaP cells at the indicated concentrations. **(E)** Immunostaining for SPINK1 and ERG using same cells as in (D). F-actin and nucleus was stained using TRITC-phalloidin and DAPI respectively. Scale bar represents 10μm. **(F)** Quantification of SPINK1 immunofluorescence images represented as relative mean fluorescence intensity (MFI) per $\mu m^2$ (in arbitrary units; A.U.) in VCaP cells treated with bicalutamide at the indicated concentrations. **(G)** Boyden chamber Matrigel invasion assay using bicalutamide (50μM) and enzalutamide (10μM) treated VCaP cells with or without androgen (R1881; 10nM) stimulation. Representative fields with the invaded cells are shown in the inset (20X magnification). **(H)** Quantitative PCR data showing relative expression of *SPINK1* in 22RV1 cells cultured in the presence androgen (8 nM DHT) for 2 months. **(I)** Same as in (H), except 22RV1 cells were cultured in the presence of bicalutamide (5μM) for 2 months. **(J)** Same as in (I), except CWR22Pc cells were used. **(K)** Same as (F), except siRNA mediated *AR* silenced VCaP cells were used (related to **Fig 2N**). **(L)** Quantitative PCR data showing relative expression of *AR* and *SPINK1* in siRNA mediated *AR* silenced 22RV1 cells with respect to control cells (NT). Immunoblot analysis for AR levels using same cells (top panel). β-actin was used as a loading control. **(M)** Immunostaining for AR and SPINK1 using same cells as in (J). F-actin and nucleus was stained using TRITC-phalloidin and DAPI respectively. Scale bar represents 10μm. **(N)** Same as (K), except siRNA mediated *AR* silenced 22RV1 cells were used.
Statistical significance was calculated by one-way ANOVA with Tukey's post hoc test for multiple comparisons in the panel (F). In all panels except (E) and (M), biologically independent samples were used (n=3); data represents mean ± SEM. *$P \leq 0.05$ and **$P \leq 0.001$ using two-tailed unpaired Student's *t* test.

#### Supplementary Figure S3

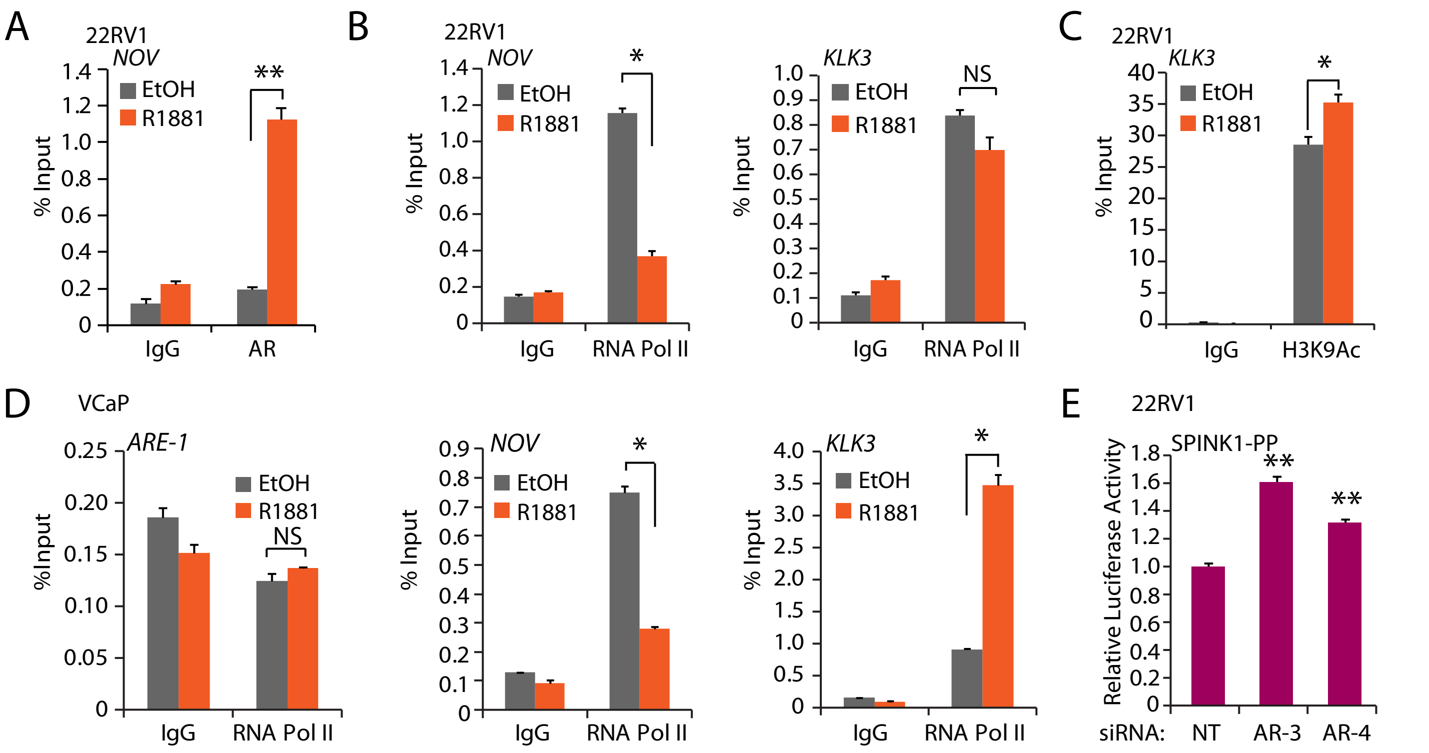


**Supplementary Figure S3. AR is recruited on *SPINK1* promoter upon androgen stimulation.**
**(A)** ChIP-qPCR data for AR occupancy on the *NOV* promoter in androgen (R1881; 10 nM) stimulated 22RV1 cells. **(B)** ChIP-qPCR data for RNA Pol II occupancy on the *NOV* and *KLK3* promoters in androgen (R1881; 10 nM) stimulated 22RV1 cells. **(C)** ChIP-qPCR data for the presence of H3 lysine 9 acetylation (H3K9Ac) marks on the *KLK3* promoter in androgen (R1881; 10 nM) stimulated 22RV1 cells. **(D)** ChIP-qPCR data for RNA Pol II occupancy *SPINK1*, *NOV* and *KLK3* promoters in androgen (R1881; 10 nM) stimulated VCaP cells. **(E)** Luciferase reporter activity of the proximal promoter of *SPINK1* (SPINK1-PP) using siRNA mediated *AR* knockdown 22RV1 cells.
For all panels, biologically independent samples were used (n=3); data represents mean ± SEM. *$P \leq 0.05$ and **$P \leq 0.001$ using two-tailed unpaired Student's *t* test.

### Supplementary Figure S4

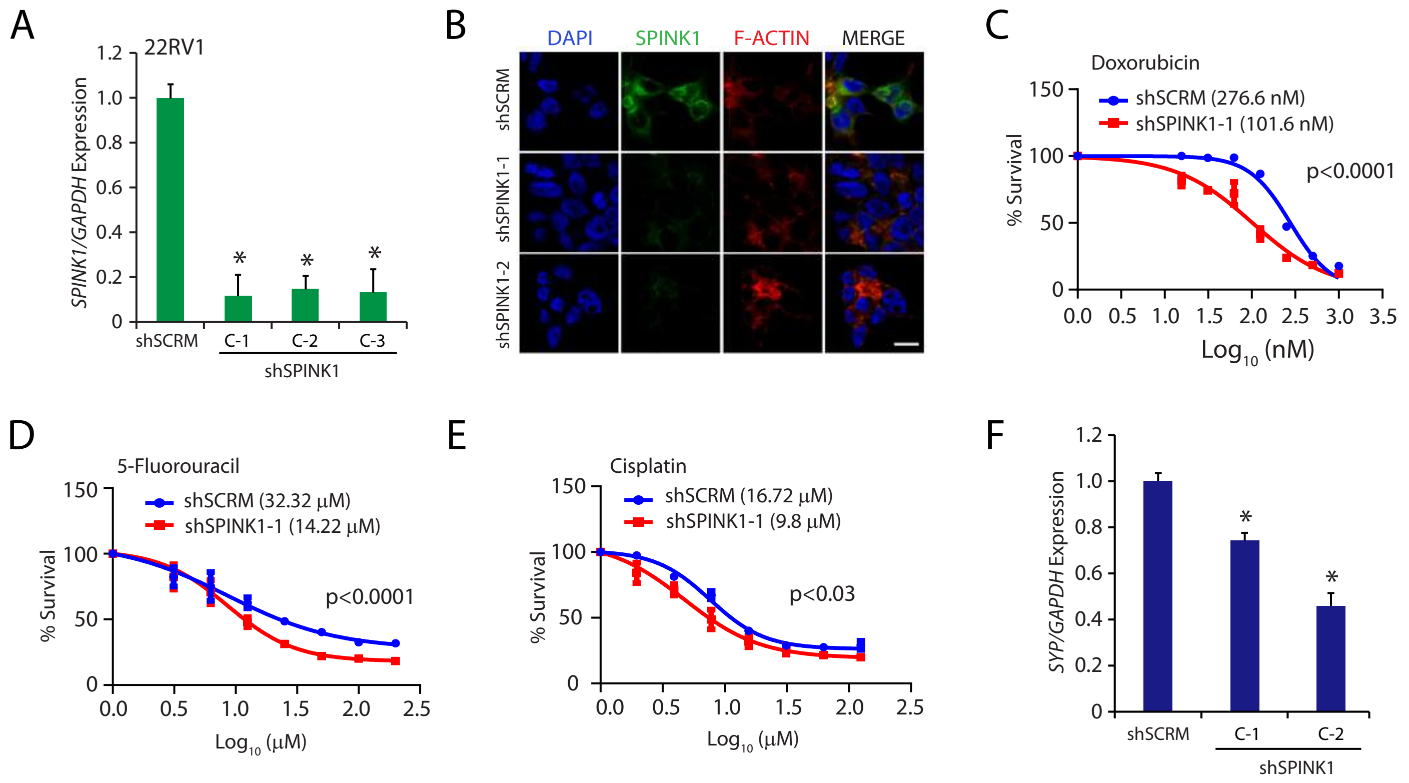


**Supplementary Figure S4. Knockdown of *SPINK1* attenuates epithelial–mesenchymal transition and stemness in PCa cells.**
**(A)** Quantitative PCR data showing relative expression of *SPINK1* in stable *SPINK1* silenced 22RV1 cells using three independent shRNA against *SPINK1* (C1, C2 and C3). **(B)** Immunostaining for SPINK1 using same cells as in (A). F-actin and nucleus was stained using TRITC-phalloidin and DAPI respectively. Scale represents 10µm. **(C)** Dose-response curve with a range of doxorubicin concentration using control 22RV1 (shSCRM) and *SPINK1* silenced (shSPINK1-1) cells. $IC_{50}$ values were calculated using GraphPad Prism software. **(D)** Same as in (C), except 5-fluorouracil was used. **(E)** Same as in (C), except cisplatin was used. **(F)** Quantitative PCR data showing relative expression of *SYP* in stable *SPINK1* silenced (shSPINK1-1) and control 22RV1 (shSCRM) cells.
For all panels except (B), biologically independent samples were used (n=3); data represents mean ± SEM. $^*P \leq 0.05$ and $^{**}P \leq 0.001$ using two-tailed unpaired Student's *t* test.

### Supplementary Figure S5

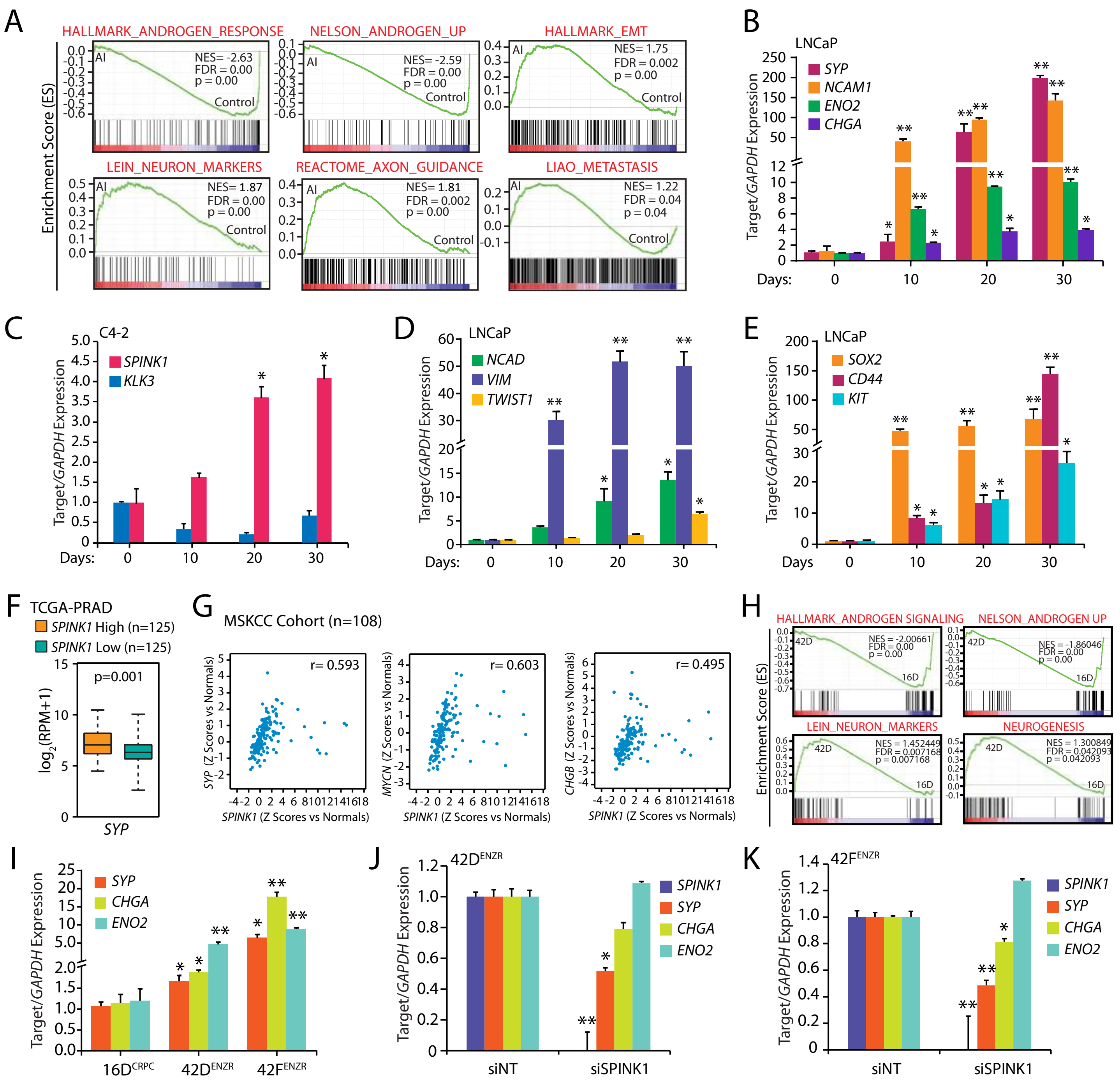

**Supplementary Figure S5. SPINK1 expression is positively correlated with neuroendocrine prostate cancer (NEPC) signature genes.**

**(A)** Gene Set Enrichment Analysis (GSEA) plots showing gene signatures associated with androgen signaling, neuroendocrine phenotype and EMT with the corresponding statistical metrics in long-term androgen deprived LNCaP cells (AI) relative to control cells (GSE8702). **(B)** Quantitative PCR data showing relative expression of *SYP*, *CHGA*, *ENO2* and *NCAM1* in long-term androgen deprived (30 days) LNCaP cells. **(C)** Same as in (B), except relative expression of *SPINK1* and *KLK3* in long-term androgen deprived C4-2B cells, a metastatic subline derived from LNCaP-C4. **(D)** Same as in (C), except relative expression of *NCAD*, *VIM* and *TWIST1*. **(E)** Same as in (C), except relative expression of *CD44*, *SOX2* and *KIT*. **(F)** Relative expression of *SYP* in SPINK1 high (n=125) and SPINK1 low (n=125) PCa patients' available at TCGA-PRAD dataset. Transcripts level shown as $\log_2$(RPM+1). *P*-values calculated using two-tailed unpaired Student's *t* test. **(G)** Correlation plots between *SPINK1* and *SYP*, *CHGB* and *MYCN* mRNA Z-scores in MSKCC PCa cohort (n=108) analyzed using cBioPortal. Spearman's correlation coefficient (r) indicated. **(H)** Gene Set Enrichment Analysis (GSEA) plots showing gene signatures associated with androgen signaling and neuroendocrine phenotype with the corresponding statistical metrics in the $42D^{ENZR}$ cells compared to $16D^{CRPC}$ cell line. **(I)** Quantitative PCR data showing relative expression of *SYP*, *CHGA* and *ENO2* in $16D^{CRPC}$, $42D^{ENZR}$ and $42F^{ENZR}$ cells. **(J)** Same as in (I), except relative expression of *SPINK1*, *SYP*, *CHGA* and *ENO2* in siRNA mediated *SPINK1* silenced $42D^{ENZR}$ cells. **(K)** Same as in (J), except $42F^{ENZR}$ cells were used.

In panels (B), (C), (D), (E), (I), (J) and (K), biologically independent samples were used (n=3); data represents mean ± SEM. *$P \leq 0.05$ and **$P \leq 0.001$ using two-tailed unpaired Student's *t* test.

### Supplementary Figure S6

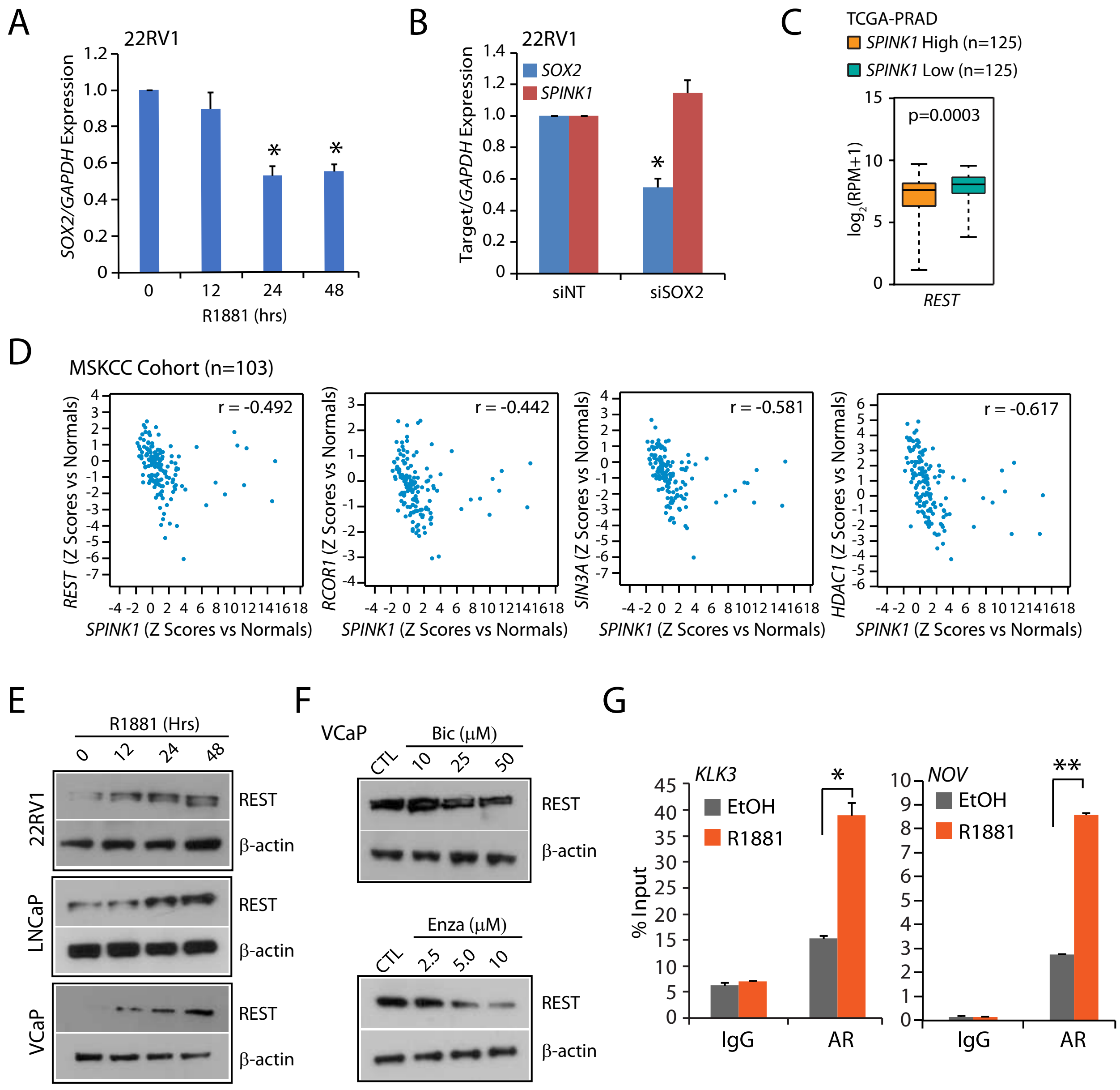


**Supplementary Figure S6. REST serves as a transcriptional co-repressor of AR and negatively regulates *SPINK1* expression.**

**(A)** Quantitative PCR data showing relative expression of *SOX2* in androgen (R1881; 10 nM) stimulated 22RV1 cells at the indicated time points. **(B)** Quantitative PCR data showing relative expression of *SOX2 and SPINK1* in siRNA mediated *SOX2* silenced 22RV1 cells with respect to control. **(C)** Relative expression of *REST* across SPINK1 high (n=125) and SPINK1 low (n=125) PCa patients' in TCGA-PRAD dataset. Transcripts level shown as $\log_2$ (RPM+1). **(D)** Correlation plots between *SPINK1* and *REST*, *RCOR1*, *SIN3A* and *HDAC1* mRNA Z-scores in MSKCC PCa cohort (n=103) analyzed using cBioPortal. Spearman's correlation coefficient (r) indicated. **(E)** Immunoblot analysis for REST level in 22RV1, LNCaP and VCaP cells upon androgen (R1881; 10 nM) stimulation at different time points as indicated. β-actin was used as a loading control. **(F)** Same as in (C), except anti-androgens (bicalutamide and enzalutamide) treated VCaP cells were used. β-actin was used as a loading control. **(G)** ChIP-qPCR data showing recruitment of AR on the *KLK3* and *NOV* promoters in androgen (R1881; 10 nM) stimulated LNCaP cells.

For panels (A), (B) and (G), biologically independent samples were used (n=3); data represents mean ± SEM. **P*≤ 0.05 and ***P*≤

#### Supplementary Methods

##### Cell lines and authentication

All the prostate cancer cell lines (22RV1, VCaP and LNCaP) were obtained from American Type Cell Culture (ATCC) and maintained as per the guidelines. Briefly, cells were cultured in recommended medium supplemented with 10% fetal bovine serum (FBS) (Gibco) and 0.5% Penicillin Streptomycin (Pen Strep) (Thermo Fisher Scientific); cultured in $CO_2$ incubator (Thermo Fisher Scientific) supplied with 5% $CO_2$ at 37°C temperature. Cell line authentication was done via short tandem repeat (STR) profiling at the Lifecode Technologies Private Limited, Bangalore and DNA Forensics Laboratory, New Delhi. Routine *Mycoplasma* contamination of all cell lines were checked using PlasmoTest mycoplasma detection kit (InvivoGen). CWR22Pc cells were a kind gift from Dr. Marja T. Nevalainen and were cultured as previously discussed (Dagvadorj et al, 2008). C4-2 cells were a generous gift from Dr. Mohammad Asim and were cultured as described previously (Asim et al, 2016). $16D^{CRPC}$, $42D^{ENZR}$ and $42F^{ENZR}$ cells were kindly gifted by Dr. Amina Zoubeidi and were cultured as previously mentioned (Bishop et al, 2017).

##### Plasmids and Constructs

pGL3-SPINK1-PP construct was obtained by cloning the *SPINK1* proximal promoter (SPINK1-PP) in pGL3-basic vector, a kind gift from Dr. Amitabha Bandyopadhyay. pGL3-SPINK1-DP was obtained by cloning distal promoter of the *SPINK1* gene in pGL3-SV40 enhancer vector (Promega). Site directed mutagenesis was performed to alter the androgen response element (ARE) in the *SPINK1* distal promoter (SPINK1-DP) to generate the pGL3-SPINK1-DP mutant (MT) from pGL3-SPINK1-DP wildtype (WT). pcDNA3.1(+) SOX2 overexpression vector was purchased from GenScript and pcDNA3.1(+) empty vector was kindly gifted by Dr. Arun K.

Shukla. Wild type and mutant (ΔNLS and V581F) AR constructs cloned in FUCGW lentiviral vectors were generously given by Dr. Owen Witte. pGIPZ plasmids (shScrambled, shSPINK1-1, shSPINK1-2 and shSPINK1-3) were procured from Dharmacon.

**Immunohistochemistry (IHC) staining**

IHC for AR and SPINK1 was performed using EnVision FLEX system (Agilent). Briefly, TMA slides were incubated at 60° C for 2 hours and antigen retrieval was done in EnVision FLEX Target Retrieval Solution, High pH (Agilent DAKO, K800421-2) in a PT Link instrument (Agilent DAKO, PT200. Slides were washed in 1X EnVision FLEX Wash Buffer (Agilent DAKO, K800721-2) for 5 minutes, followed by treatment with Peroxidazed 1 (Biocare Medical, PX968M) for 5 minutes and Background Punisher (Biocare Medical, BP974L) for 10 minutes with a wash after each step. Mouse monoclonal SPINK1 (Novus Biologicals, H00006690-M01, 1:100 dilution) and AR (CST, 5153, 1:200 dilution) antibody diluted in EnVision FLEX Antibody Diluent (Agilent DAKO, K800621-2) was added to each slide and incubated overnight at 4º C. Slides were washed and incubated in Mach2 Doublestain 1 (Biocare Medical, MRCT523L) for 30 minutes at room temperature. Subsequently, slides were rinsed in 1X EnVision Wash Buffer thrice and then treated with a Ferangi Blue solution (Biocare Medical, FB813S) for 7 minutes. Slides were rinsed twice in distilled water and stained with EnVision FLEX Hematoxylin (Agilent DAKO, K800821-2) for 5 minutes. After multiple washes, slides were immersed in a 0.01% ammonium hydroxide solution and rinsed twice in distilled water. Once the slides were dried completely, they were put in xylene for approximately 15 times and ultimately mounted using EcoMount (Biocare Medical, EM897L).

**Analysis of TCGA-PRAD data**

For understanding the inverse association between SPINK1 and AR signaling in prostate cancer patients, Illumina HiSeq mRNA data of patients with prostate adenocarcinoma (PRAD) was downloaded from TCGA portal for *SPINK1*, *AR*, *NCOR1*, *NCOR2*, *NRIP1*, *SYP* and *REST* genes. Since, *SPINK1* gene is overexpressed in ~10-15% of the PCa patients (Tomlins et al, 2008), therefore we performed quartile based normalization (Dillies et al, 2013) to stratify the patients based on high and low *SPINK1* expression. Accordingly, patients corresponding in the top quartile (n=125) (QU, $\log_2$ (RPM+1)>5.468 or $\log_2$ (normalized count+1)>1.892) were considered as *SPINK1*-high whereas patients in the lower quartile (n=125) (QL, $\log_2$ (RPM+1)<1.124 or $\log_2$ (normalized count+1)<-2.611) were assigned as *SPINK1*-low The corresponding gene expression values for *AR*, *NCOR1*, *NCOR2*, *NRIP1*, *SYP* and *REST* in *SPINK1*-high versus *SPINK1*-low patients were compared to identify the association between *SPINK1* and these genes. For the heatmap between *AR* and *SPINK1*, PCa patients from TCGA-PRAD cohort were grouped based on high and low *AR* expression ($\log_2$ (RPM+1) >9.4 and $\log_2$ (RPM+1) <8.1). High and low cutoff value for *AR* expression was obtained from the TCGA portal. The corresponding *SPINK1* expression values in *AR*-high and *AR*-low patients was further used to construct the heatmap by employing 'gplot' function in 'R'. Correlation plots between *SPINK1* and *REST* and its complex members derived from the MSKCC cohort were directly retrieved from cBioPortal (http://www.cbioportal.org/).

**Lentiviral Packaging**

Lentiviral particles were produced using ViraPower Lentiviral Packaging Mix (Invitrogen) according to the manufacturer's instructions. Briefly, HEK293FT were plated at 90% confluency in a 100 mm culture dish, in antibiotic free media and transfected with the plasmids of the packaging mix along with pGIPZ plasmids (9µg+3µg) using FuGENE HD Transfection Reagent

(Promega). 24 hours post transfection, the media was replaced, and the viral particles were harvested 48-60 hours later by collecting the media. Viral particles were aliquoted and stored at -80°C to prevent repeated freeze-thaw. For generating stable lines, 22RV1 and LNCaP cells were plated in a 6-well dish and infected with viral particles along with polybrene (hexadimethrine bromide; 8µg/ml) (Sigma-Aldrich). Next day, the media was replaced and three days of post-infection, cells were selected in puromycin (Sigma-Aldrich, R0908) at a concentration of 1µg/ml.

**Androgen stimulation and deprivation**

For androgen stimulation, cells were first starved for 72 hours in phenol red free (PRF) medium supplemented with 5% charcoal stripped serum (CSS) (Gibco) and then stimulated with synthetic androgen, 10nM R1881 (Sigma-Aldrich) for stipulated period. For anti-androgen treatment, VCaP cells were serum starved for 8 hours in DMEM medium supplemented with GlutaMAX (Gibco) and then treated with different concentrations of enzalutamide (MedChemExpress, HY-70002) and bicalutamide (Sigma-Aldrich, B9061) for 48 hours in complete media. For long-term androgen deprivation, LNCaP cells were cultured in RPMI-1640 phenol red free medium (Gibco) supplemented with 5% CSS for a duration of 30 days.

**Casein kinase 1 inhibitor (iCK1) treatment**

22RV1 cells were serum starved for 12 hours in RPMI-1640 medium (Gibco) and then treated with different concentrations of iCK1, D4476 (MedChemExpress, HY-10324) for 60 hours in complete media.

**Transient Transfection**

22RV1 and VCaP cells were plated at 40-45% confluency and transfected with 30pmol of small interfering RNA (siRNA) against *AR* (Dharmacon, Cat No. LU-003400-00-0002), *SPINK1* (Dharmacon, Cat. No. LU-019724-00-0002), *SOX2* (Thermo Fisher Scientific, Cat No. 4392420) and non-targeting control (Dharmacon, Cat. No. D-001810-10-05) using Lipofectamine RNAiMAX Transfection Reagent (Thermo-Fisher Scientific) according to manufacturer's instructions. The cells were again transfected after 24 hours, and harvested for RNA and protein expression analysis, 36 hours post second transfection.

**Real Time Quantitative PCR**

Total RNA was extracted using TRIzol (Ambion) and cDNA synthesis of 1µg of RNA was performed using SuperScript III First-Strand Synthesis System (Invitrogen) according to the manufacturer's instructions. For quantitative PCR (qPCR), all reactions were performed in triplicates using SYBR Green PCR Master Mix (Applied Biosystems). Relative target gene expression was calculated for each sample using the ΔΔCt method as described before (Ateeq et al, 2011), using primers mentioned in the Supplementary Table S2.

**Gene expression array analysis**

For gene expression profiling, the total RNA from 22RV1 SCRM, shSPINK1-1, shSPINK1-2 and shSPINK1-3 cells was collected and subjected to Agilent Whole Human Genome Oligo Microarray profiling (dual color) according to manufacturer's protocol using Agilent platform (8x60K format). Three separate microarray hybridizations were performed, using 22RV1 shSPINK1 cells against control SCRM cells. For the microarray data 'Lowess' (locally weighted regression) normalization was performed. The gene expression pattern for differentially regulated genes was identified using hierarchical clustering implemented Pearson coefficient correlation algorithm. Benjamini and Hochberg procedure was used to calculate FDR-corrected *P*-values (with FDR< 0.05) to identify differentially expressed genes. Differentially expressed genes ($\log_2$ fold change > 0.5 or < -0.5, $P< 0.05$) were then used for identifying the pathways using Database for Annotation, Visualization and Integrated Discovery (DAVID) (Huang da et al, 2009a; Huang da et al, 2009b). Gene set enrichment analysis (GSEA) was performed to analyze publicly available GEO dataset (GSE8702) to identify gene-sets enriched in androgen deprived LNCaP cells. RNA-sequencing (RNA-Seq) data of $42D^{ENZR}$ and $16D^{CRPC}$ cells was obtained from Dr. Amina Zoubeidi's lab (Bishop et al, 2017). Heatmap.2 function of 'gplots' in R was used to make heatmaps.

**Western Blot Analysis**

Cell lysates were prepared in radioimmunoprecipitation assay (RIPA) lysis buffer, along with cOmplete Protease Inhibitor Cocktail (Roche) and Phosphatase Inhibitor Cocktail Set II (Calbiochem). Protein samples subjected to SDS-PAGE were transferred onto a polyvinylidene difluoride (PVDF) membrane (GE Healthcare). The membrane was blocked with 5% non-fat dry milk in Tris-buffered saline, 0.1% Tween 20 (TBS-T) for one hour at room temperature, and then incubated overnight at 4ºC with the following primary antibodies: 1:1000 diluted anti-AR (CST, 5153), 1:2000 diluted anti-REST (Abcam, ab75785), 1:2000 diluted anti-PSA (CST, 5877), 1:2000 diluted anti-SYP (Abcam, ab32127), 1:1000 diluted anti-SOX2 (Abcam, ab97959) and 1:5000 diluted anti-β-Actin (Abcam, ab6276). Subsequently, blots were washed in TBS-T and incubated with respective horseradish peroxidase-conjugated secondary anti-mouse or anti-rabbit antibody (Jackson ImmunoResearch) for 2 hours at room temperature. After washing, the signals were visualized by enhanced chemiluminescence system as described by the manufacturer (GE Healthcare).

**Chromatin Immunoprecipitation Assay**

Cells were crosslinked for 10 mins with 1% formaldehyde followed by quenching with 125mM Glycine for 5-8 mins at room temperature. The cells were washed twice with chilled phosphate buffered saline (1X PBS, pH 7.4) and lysed with the lysis buffer [1% SDS, 50mM Tris-

Cl (pH 8.0), 10mM EDTA and protease inhibitor cocktail (Roche)]. The cell lysate was then sonicated using Bioruptor (Diagenode) to obtain DNA fragments of length ~500bp. The sheared chromatin was collected after centrifugation and incubated overnight at 4°C with 4µg of either primary or isotype control antibodies. Chromatin immunoprecipitation was carried out using the following antibodies: AR (CST, 5153), Rpb1 CTD (CST, 2629), H3K9Ac (CST, 9649), REST (Abcam, ab70300), SOX2 (Abcam, ab97959) and isotype control antibodies, rabbit IgG (Invitrogen, Cat No. 10500C) and mouse IgG (Invitrogen, Cat No. 10400C). Simultaneously, Dynabeads coated with Protein G (Invitrogen) were blocked using 500µg/ml of sheared salmon sperm DNA (Sigma-Aldrich) and 100µg/ml bovine serum albumin (BSA) (HiMedia) overnight at 4°C. Blocked beads were washed using dilution buffer [1% Triton X-100, 2mM EDTA, 150mM NaCl, 20mM Tris-Cl (pH 8.0) with protease inhibitor cocktail (Roche)] and then incubated for 6-8 hours at 4°C with the lysate containing antibody to make antibody-bead conjugates. After the incubation, the beads conjugated with antibody were washed thrice in low salt wash buffer (1% Triton X-100, 150mM NaCl, 0.1% SDS, 20mM Tris-HCl (pH 8.0), 2mM EDTA with protease inhibitor cocktail) and once with high salt wash buffer (same as previous wash buffer except 500mM NaCl) followed by a final wash with 1X TE buffer. The immunocomplex was then eluted using elution buffer [1% SDS, 100mM $NaHCO_3$, Proteinase K (Sigma-Aldrich) and RNase A (500µg/ml each) (Sigma-Aldrich)]. DNA isolation was done using phenol-chloroform-isoamyl alcohol extraction. Precipitated DNA was washed with 70% ethanol, air dried and resuspended in nuclease free water (Ambion). The ChIP-qPCR was performed using primers mentioned in the Supplementary Table S2.

**Chromatin Immunoprecipitation Sequencing (ChIP-Seq) data analysis**

To determine the recruitment of AR on the *SPINK1* promoter, we analyzed publicly available ChIP-Seq data (GSE58428) for AR pull down in VCaP cells, treated with DHT and vehicle control, ethanol (EtOH). Raw single-end reads were first analyzed for their quality using FASTQC, followed by trimming with FASTQ Trimmer, ensuing all the default settings of Galaxy web platform available on the public server at usegalaxy.org (Afgan et al, 2018). Reads were aligned to the reference genome (hg18) using Bowtie to generate Sequence Alignment/Map (SAM) files. Unaligned or unmapped reads were filtered using FilterSAM, a utility of SAMtools. These SAM files were converted to its Binary Alignment/Maps (BAM) files using SAMtools. Further, ChIP-Seq peaks for DHT and EtOH treated samples were called using Model-based analysis of ChIP-seq (MACS; $P < 10^{-5}$) with default settings against Input. BAM and BED files obtained were visualized by Integrative Genomic Browser (IGB) (Freese et al, 2016).

**Immunofluorescence**

Cells were grown on glass coverslips in 24-well culture dishes and fixed with 4% paraformaldehyde (in 1X PBS). The cells were washed with 1X PBS, permeabilized with 0.3% Triton X-100 in 1X PBS (PBS-T) for 10 mins and then blocked using 5% normal goat serum in PBS-T for 2 hours at room temperature. The cells were then incubated with primary antibodies: SPINK1 (1:100, Abnova H00006690-M01), AR (1:200, CST 5153), ERG (1:200, ab92513), E-cadherin (1:400, CST 3195), Vimentin (1:100, CST 3932) diluted in PBS-T, overnight at 4ºC. Cells were then washed using 0.05% Tween 20 in 1X PBS, followed by incubation with Alexa Fluor conjugated anti-mouse or anti-rabbit secondary antibodies (1:600 dilution, CST #4408 and #4412, respectively). The cells were washed again and stained with TRITC-Phalloidin (Sigma-Aldrich) followed by DAPI (Sigma-Aldrich). The coverslips were mounted on glass slides using Vectashield mounting medium (Vector laboratories). Stained cells were visualized, and images were captured using Axio Observer Z1 inverted fluorescence microscope (Carl Zeiss) equipped with an Apotome device. For the quantification of SPINK1 expression, stained cells were visualized, and images were captured with a 60X oil immersion objective (NA 1.4). Post-processing of the acquired images was done using ImageJ software. The boundary of each cell was marked and the cells with area ranging between 70-130 $\mu m^2$ were considered to quantify mean fluorescence intensity per unit area for each condition. Statistical significance was calculated using one-way ANOVA or t-test depending on the number of groups in the datasets. For neurite outgrowth quantification, 20 random fields were imaged for stable LNCaP shSPINK1 green fluorescent protein (GFP) positive cells cultured in the androgen deprived condition for a duration of 30 days. The neurite lengths were measured by using the Simple Neurite Tracer as discussed previously (Longair et al, 2011).

**Luciferase Promoter Reporter Assay**

22RV1 cells were plated at 40-50% confluency in a 24-well plate and transfected with 250ng of pGL3-SPINK1-PP along with 2.5ng of pRL-null vector (Promega), an internal control, using FuGENE HD Transfection Reagent (Promega). After 24 hours of transfection, cells were starved in RPMI-1640 phenol red free (PRF) medium (Gibco) for 8 hours and stimulated with synthetic androgen, 10nM R1881 (Sigma-Aldrich) in RPMI-PRF medium containing 5% charcoal stripped serum (CSS) for 18 hours. Cells were harvested using the lysis buffer provided with Dual-Glo Luciferase assay kit (Promega). Firefly and Renilla luciferase activity were measured according to the manufacturer's protocol. For each sample, firefly luciferase activity was normalized to Renilla luciferase activity. Similar protocol was followed to measure the luciferase promoter reporter activity for pGL3-SPINK1-DP WT, pGL3-SPINK1-DP- MT and pGL4.10-PSA constructs.

**Prostatosphere Assay**

22RV1 SCRM and shSPINK1 cells ($1\times10^4$) were plated in low adherence 6-well cell culture dishes in serum-free DMEM-F12 (1:1, Invitrogen) supplemented with B27 (50X, Invitrogen), EGF (20 ng/ml, Invitrogen), FGF (20 ng/ml, Invitrogen) as previously described (Dontu et al, 2003). A small population of cells forming prostatospheres were collected by centrifugation and dissociated into cell suspension of single cells. The cells were passaged in the similar manner for several generations and the experiment was terminated after two weeks. The prostatospheres formed were assessed for sphere forming efficiency. Mean area of spheres were measured using ImageJ software and spheres >50µm in diameter were counted and the values were represented as percent sphere formation efficiency. The three-dimensional (3D) tumor spheroid assay was performed as described previously (Pal & Kleer, 2014). Briefly, each well of chamber slide was coated with growth factor reduced (GFR) Matrigel (Corning) and incubated at 37$^{o}$C. 22RV1 cells ($3\times10^3$) were resuspended in complete media supplemented with 2% Matrigel and plated in the Matrigel precoated chamber slide. Complete media supplemented with Casein Kinase 1 inhibitor, iCK1 (20µM) or DMSO control was changed after every two days until 2 weeks. The tumor spheroids of size >1000µm in diameter were counted and the values were represented as percent sphere formation efficiency and the mean area of spheres were plotted.

**Migration Assay**

Migration assay was performed using Transwell Boyden chamber of 8µm pore size (Corning). Briefly, after the desired treatment, cells were trypsinized, counted and $1\text{x}10^5$ cells were resuspended in serum free DMEM media supplemented with GlutaMAX (Gibco) and added to the upper chamber of the Transwell. The bottom chamber was supplemented with RPMI-1640 media supplemented with 30% FBS. After 48 hours of incubation, the migrated cells were fixed with 4%

paraformaldehyde (in 1X PBS) and stained with 0.5% (w/v) crystal violet. Representative pictures were captured using Axio Observer Z1 microscope (Carl Zeiss). For quantification, the Transwells were destained using 10% acetic acid (v/v) in distilled water. Absorbance was measured at 550 nm.

**$IC_{50}$ determination**

For determining the $IC_{50}$ of drugs, 22RV1 SCRM and shSPINK1-1 cells ($3x10^3$) were plated in 96-well dishes and treated with varying concentration of drugs for 48 hours. The $IC_{50}$ of the drugs was determined using WST-1 (Roche) as described previously (Arriaga et al, 2014).

**Side population (SP) Assay**

22RV1 SCRM and shSPINK1 cells ($5x10^5$) were stained with Hoechst 33342 (5μg/ml) (Thermo Fisher Scientific) with agitation for 2 hours at 37 °C in a water bath. The Hoechst staining was performed in absence and presence of verapamil (Sigma-Aldrich), ATP-binding cassette transporter (ABC transporter) inhibitor. The cells were then washed and resuspended in FACS buffer (1X PBS, 2% fetal bovine serum and 0.1% sodium azide) and subjected to flow cytometry. To exclude the dead cells, Propidium iodide (PI) (5μg/ml) (BioLegend) staining was performed. The side population (SP) was detected using UV laser at 350 nm. The blue fluorescence was captured using 460/50 band-pass filter while the red fluorescence was measured using 670/30 long pass filter. Higher laser power was used to capture red channel as its intensity is lower than blue channel. Optimal resolution of SP cells was obtained by using a laser power of 30-35 mW. Cells were first gated for the live population; SP tails were then gated and displayed as a dot plot for Hoechst blue and red scatter. For every condition, ~$1x10^5$ events were acquired. Data analysis and acquisition was done using the FlowJo software.

**ALDH assay**

To determine the enzymatic activity of aldehyde dehydrogenase (ALDH), Aldefluor assay was performed using Aldefluor kit (Stem Cell Technologies) following the manufacturer's guidelines. Briefly, 22RV1 cells were transfected twice with si*SPINK1* or control siRNA (si*NT*) at an interval of 24 hours. After 36 hours of second transfection, the cells were collected, washed twice with 1X PBS and harvested by incubating with cell dissociation solution, Cellstripper (Corning). The cells were then washed using 1X PBS and centrifuged at 1500 rpm for 5 mins at 4°C. Afterwards, the cells were resuspended in 1ml Aldefluor assay buffer followed by addition of 5µL of activated Aldefluor substrate. Cell suspension was immediately divided in two parts, one tube having 5µL of ALDH inhibitor, diethylaminobenzaldehyde (DEAB) served as a negative control, and the other tube was used as such. The cells were incubated at 37°C for 30 mins, centrifuged and resuspended in 500µL of Aldefluor assay buffer. The Aldefluor activity was detected in green channel. The cells treated with DEAB served as a respective control for gating the ALDH+ cell population. The ALDH activity of cells transfected with si*SPINK1* were compared with the cells transfected with si*NT*. For data acquisition, BD FACSCanto II (BD Biosciences) was used and the analysis was performed using FlowJo software.

**Flow Cytometry**

22RV1 cells transfected with siRNAs, si*SPINK1* and si*NT*, were washed with 1X PBS and collected using cell dissociation solution, Cellstripper (Corning). Staining with anti-human NCAM1 antibody (eBioscience, Ab 25-0567-42) was performed by incubating the cells with the antibody (1:50 dilution) for one hour at 4°C, followed by staining with 7-Aminoactinomycin D (7-AAD) for cell viability. Data acquisition was done using BD FACSCanto II (BD Biosciences) and analyzed with FlowJo software.

**Cell viability assay**

Cell viability assay was carried out by plating 3000 cells per well in 96-well plate. Cells were treated with the indicated drug concentrations (10µM and 20µM) against DMSO control and incubated upto 48 hours. Cell viability was determined following incubation with the Cell Proliferation Reagent WST-1 (Roche), followed by colorimetric assay as per manufacturers protocol.

**Foci formation assay**

22RV1 cells ($2\times10^3$) were plated in six-well culture dishes in RPMI-1640 medium (Gibco) supplemented with 5% heat-inactivated fetal bovine serum (Invitrogen) and incubated at 37°C, media was changed after every 48 hours with Casein Kinase 1 inhibitor, iCK1 (20µM) along with DMSO control. The assay was terminated after 2 weeks and foci were fixed in paraformaldehyde (4% in PBS) and stained with crystal violet solution (0.1% w/v). For destaining, 10% glacial acetic acid was used, and the absorption was quantified at 550nm.
